# A theoretical basis for cell deaths

**DOI:** 10.1101/2024.03.04.583348

**Authors:** Yusuke Himeoka, Shuhei A. Horiguchi, Tetsuya J. Kobayashi

## Abstract

Understanding deaths and life-death boundaries of cells is a fundamental challenge in biological sciences. In this study, we present a theoretical framework for investigating cell death. We conceptualize cell death as a controllability problem within dynamical systems, and compute the life-death boundary through the development of “stoichiometric rays”. This method utilizes enzyme activity as control parameters, exploiting the inherent property of enzymes to enhance reaction rates without affecting thermodynamic potentials. This approach facilitates the efficient evaluation of the global controllability of models. We demonstrate the utility of our framework using its application to a toy metabolic model, where we delineate the life-death boundary. The formulation of cell death through mathematical principles provides a foundation for the theoretical study of cellular mortality.

**SIGNIFICANCE STATEMENT:** What is death? This fundamental question in biology lacks a clear theoretical framework despite numerous experimental studies. In this study, we present a new way to understand cell death by looking at how cells can or cannot control their states. We define a “dead state” as a state from which a cell cannot return to being alive. Our method, called “Stoichiometric Rays”, helps determine if a cell’s state is dead based on enzymatic reactions. By using this method, we can quantify the life-death boundary in metabolic models. The present framework provides a theoretical basis and a tool for understanding cell death.

## I. INTRODUCTION

Death is one of the most fundamental phenomena, and comprehension of the life-death boundary is a pivotal issue in the study of biological systems. Cell death within simple unicellular model organisms such as *Escherichia coli* and yeast has been extensively studied [1–12].

What do we call “death” and from what features of a given cell do we judge the cell is dead? Despite extensive studies, such characterization of cell death is still under debate [11, 13–17]. Indeed, there is a debate as to whether the viable but non-culturable (VBNC) state [18, 19] is a dead state or not, without a clear definition of death [20, 21]. Confusion due to lack of a clear definition has also occurred in research on bacterial persistence [22–25] and the community has reached a consensus on the definition of persister [26] to avoid unnecessary slowing down of research. A clear definition of death is essential for the advancement of cell death research.

The current circumstances of microbial cell death research are in stark contrast to other topics in systems biology, where both experimental and theoretical approaches are integrated to explore cellular phenomena quantitatively and to elucidate the underlying biochemical design principles. Microbial adaptation, robustness of living systems, and network motifs are hallmarks of crosstalk between experimental and theoretical approaches [27–31]. However, despite substantial experimental data, theoretical explorations of cell death remain underdeveloped. Theoretical studies on microbial death have primarily focused on estimating death rates and assessing their impacts on population dynamics supposing a predefined concept of “death” [5–8, 10]. Theories for delineating what is death and evaluating cellular viability are indispensable for extracting the quantitative nature of cell death, and accordingly, life. The necessity for theories of cell death is beyond comprehending experimental observations; it is imperative for the advancement of biological sciences, particularly through the integration of burgeoning computational technologies. Recent developments in computational biology have facilitated the construction of whole-scale kinetic cell models [32] and the application of advanced machine-learning technologies, enabling large-scale simulations of cellular responses to environmental and genetic changes [33, 34]. This includes modeling scenarios for cell death as well as cell growth [32, 35]. The development of a robust mathematical framework to define and evaluate cell death within these models is essential to advance our understanding of cellular mortality.

In the present manuscript, we aimed at constructing a theoretical foundation for cell death using cell models described by ordinary differential equations. We introduce the definition of the dead state based on the controllability of the cellular state to predefined living states. Next, we develop a tool for judging whether a given state is dead for enzymatic reaction models, the *stoichiometric rays*. The method leverages the inherent feature of enzymatic reactions in which the catalyst enhances the reaction rate without altering the equilibrium. The stoichiometric rays enables the efficient computation of the global controllability of the models.

## II. A FRAMEWORK FOR CELL DEATHS

In the present manuscript, we formulate the death judgment problem as a controllability problem of mathematical models of biochemical systems. Figure 1 is a graphical summary of the framework proposed in the present manuscript. Herein, we focus on well-mixed deterministic biochemical reaction systems. The dynamics of these systems are described by the following ordinary differential equations:

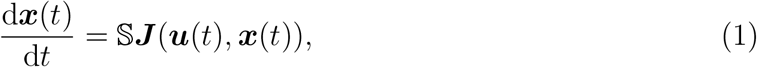

where 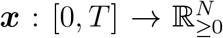, ***u*** : [0, *T*] → ℝ^*P*^, and 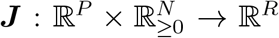 represent the concen-tration of the chemical species, control parameters, and reaction flux, respectively. Here we denote the number of chemical species, control parameters, and reactions as *N, P* and *R*, respectively, and *S* is *N* × *R* stoichiometric matrix.

**FIG. 1.**
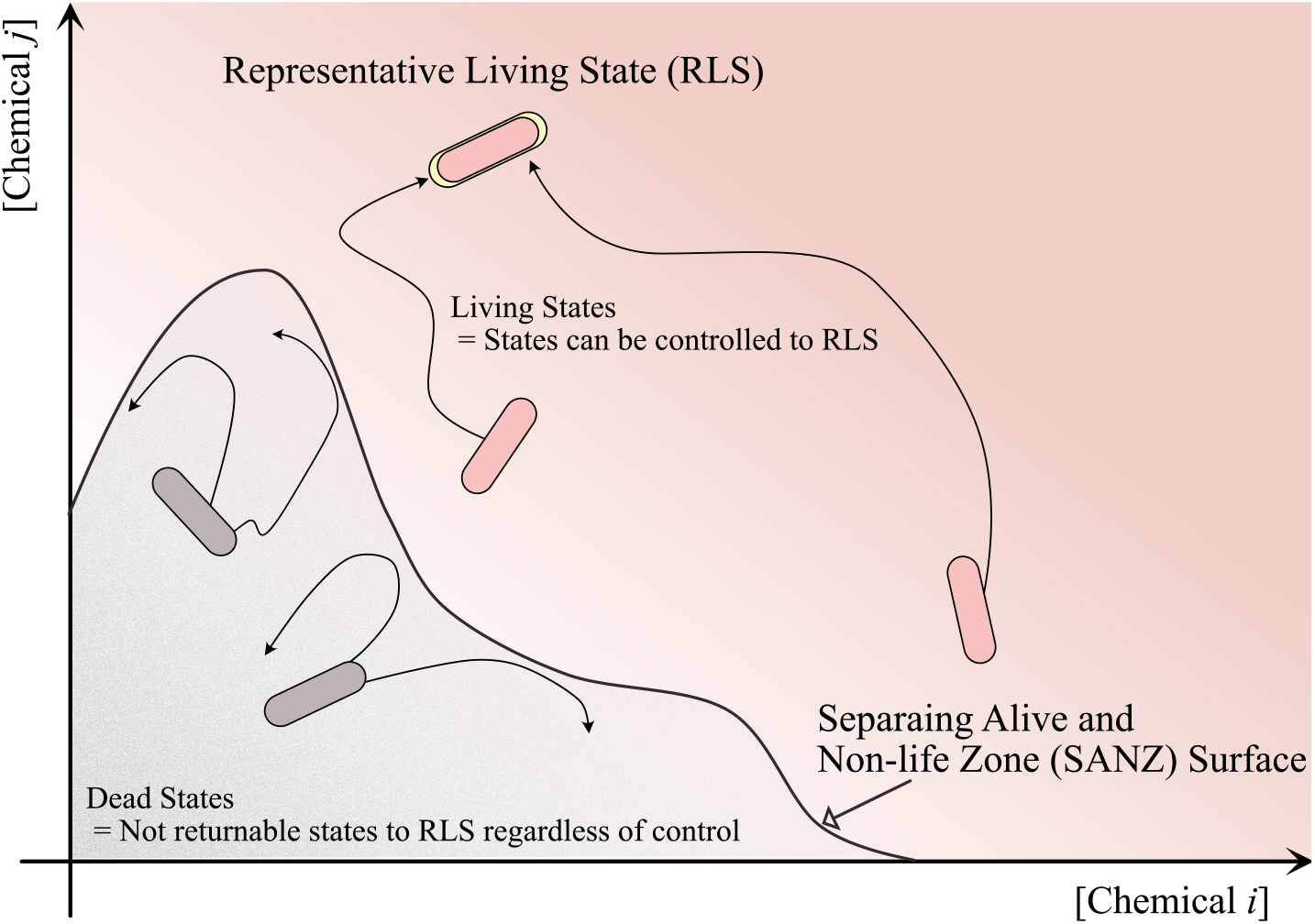
A graphical summary of the proposed framework. We consider “death” as the loss of controllability to the predefined representative living states (RLS), *X*. If a given state can be controllabled back to the RLS through modulations of biochemical parameters (concisely defined in the main body), the state is judged to be “living”. If no control back to the RLS is possible, regardless of the control, the state is “dead” in our framework. We call the set of dead states the dead set *D*(*X*) as a function of the RLS. The boundary between the controllable set and the dead set *∂C*(*X*) ∩ *∂D*(*X*) is called *Separating Alive and Non-life Zone (SANZ) hypersurface* Γ(*X*), where SANZ is taken from *Sanzu River*, the mythological river that separates the world of life and the afterlife in Japanese Buddhism [36]. In the figure, the pink and gray colours represent the living and dead states respectively. The RLS is by definition living and is coloured pink with the yellow highlighting. The two states in the middle right are considered living and are coloured pink because they can be controlled back to RLS. On the other hand, the two states in the left middle are coloured gray because they cannot be controlled back to RLS. The existence of control back to the RLS can be computed by the proposed method *the stoichiometric rays* (Sec. III) for models where the enzymatic activities and external metabolites concentrations as the control parameters. Note that our framework does not say anything about which states should be in the RLS, while some useful mathematical arguments for the choice of the RLS are provided in the Appendix.

In the following, we formally introduce the definition of dead states for the further mathematical arguments in the appendix. Intuitively, we first need to define the “representative living states” which are reference points of the living states. The choice of the representative living state is not given by the framework. We assume that one has a working hypothesis about which states should be chosen as representative living states. Then we define the dead states as the state from which it is impossible to return to any of the representative living states, regardless of control, such as modulation of enzymatic activities and the external concentrations of metabolites.

### Definition 1

(Trajectories). *We gather all the solutions of Eq*. (1) *satisfying* ***x***(0) = ***x***_0_ *and* ***x***(*T*) = ***x***_1_ *and denote the set by* 𝒯(*T*, ***x***_0_, ***x***_1_). *In addition, the union of* 𝒯 (*T*, ***x***_0_, ***x***_1_) *for all T* ≥ 0 *is denoted as*

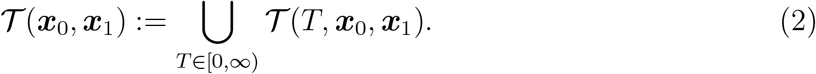

Note that 𝒯(*T*, ***x***_0_, ***x***_1_) and 𝒯(***x***_0_, ***x***_1_) contain all the solutions obtained by varying the control parameter ***u***(*t*).

### Definition 2

(Controllable Set). *The controllable set of given state* ***x*** *is defined as the set of states from which the state can be controlled to the target state* ***x***. *It is given by*

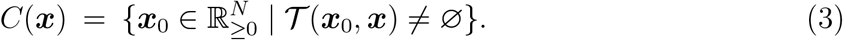

*We define the controllable set of a given set X as a union of the controllable set of points in X, that is*,

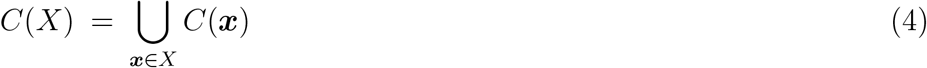

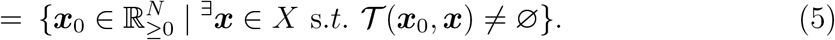

In the present framework, we assume that the *representative living states X* are given a priori as reference points of the living states. Using the controllable set and representative living states, we define the dead state as follows:

### Definition 3

(Dead State). *A state* ***z*** *is called the dead state with respect to the representative living states X if*

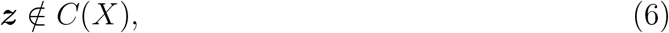

*holds*.

*The set of all dead states with respect to X form the dead set and is denoted as D*(*X*), *i*.*e*.,

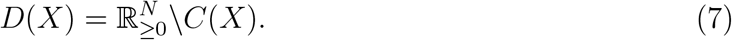

*We term the boundary of the controllable set and the dead set Γ*(*X*) ≔ ∂*C*(*X*) ∩ ∂*D*(*X*) *as the Separating Alive and Non-life Zone (SANZ) hypersurface*.

The *SANZ hypersurface* is derived from a mythical river in the Japanese Buddhist tradition, the Sanzu River, which separates the world of the living and the afterlife [36].

Note that the “death” is defined here based sorely upon the controllability of the state. The dead state can be a point attractor, limit cycle, infinitely long relaxation process, and etc. As Def. 3 states nothing on the features of the dead state except the controllability, the dead states can have, for instance, metabolic activity comparable to that of the representative living states. In such cases, the “dying state” would be an appropriate word to describe the state, while we use the term “dead state” for uniformity ^1^.

In the context of microbiology, for example, the steadily growing states are a straightforward choice of representative living states, while it is not necessary to be the growing states. In this case, the definition of death (Def. 3) can be understood by analogy with the plating assay. In the plating assay, cells are placed on the agar plate and are considered alive if they can form a colony within a given time period. The present definition with the growth state as the representative living state checks the potential for regrowth. In this sense, the definition is parallel to the plating assay. The differences lie in the universality of the controls that the cells can perform and the time scale of the control. In the present framework, we assume that cells are universal chemical systems that are able to find and execute a control to restart growth if there is a control to do so. We also do not care about the time scale of the control. The cells are considered to be living if they can regrow even if it takes an infinite amount of time, i.e. our “agar plate” never dry out.

It is also noteworthy that if the continuously growing states are chosen as representative living states, the quiescent states such as the dormant states, spore and so on should be judged as living states, since cells in these states typically retain potentials for regrowth.

The dead state and the dead set depend on the choice of representative living states. We have presented useful mathematical statements for choosing the representative living states in the Appendix. Readers interested in the choice of representative living states are encouraged to read that section, although it is not necessary for understanding the main body of this manuscript.

## III. THE STOICHIOMETRIC RAYS: A TOOL FOR THE DEATH JUDGMENT

Thus far, we have proposed a possible definition of the dead state. This definition raises the question of whether such global controllability can be computed. In this section, we show that the controllable set is efficiently computable for models with enzymatic activity and the external concentration of chemicals as the control parameter. Before moving to the main body, we briefly describe previous studies related to the controllability of chemical reaction systems.

Most biochemical reactions can take place within reasonable timescales only by the catalytic aid of enzymes [37], and thereby, the control of intracellular states relies on the modulation of enzyme activities and concentrations (hereafter collectively referred to as activities). Thus, it is indispendable to develop a mathematical technique to compute the controllability of intracellular states by modulating enzyme activities for constructing a framework of cellular viability in terms of Def. 3.

Generic frameworks for the controllability of chemical reaction systems have been studied in the control theory field [38–41]. However, there are critical limitations in previous studies when applied to biological systems; Saperstone *et al*. [38] studied the controllability of linear reaction rate functions, with which we could not model many-body reactions. In Farkas *et al*. [39] and Dochain *et al*. [40], nonlinear models were dealt with, but only local controllability was discussed. In contrast, global controllability of a nonlinear model was studied by Drexler *et al*. [41], although they allowed the kinetic rate constants to be negative-valued, to which no physical reality corresponds.

In the following, we present a simple method to compute the global controllability of enzymatic reaction models, the *stoichiometric rays*. The concept is hinted at the inherent feature of enzymatic reactions; enzymes enhance reactions without altering the equilibrium [42]. We deal with the phase space to the positive orthant, 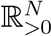^2^.

Hereafter, we restrict our attention to the cases in which the reaction rate function in Eq, (1) is expressed by the following form

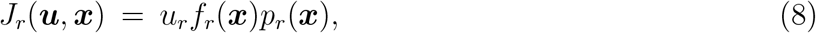

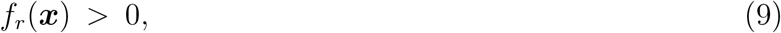

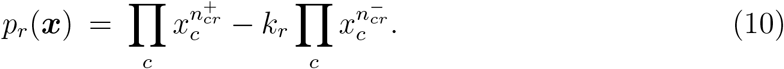

We refer to ***f*** (***x***) and ***p***(***x***) as the kinetic, and thermodynamic parts, respectively. f_*r*_(***x***) is a strictly positive function. 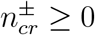 represents the reaction order of chemical c in the forward (+) or backward (−) reactions of the rth reaction. k_*r*_ ≥ 0 corresponds to the Boltzmann factor of the reaction *r*. 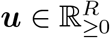 is the control parameter, representing the activities of the enzymes. In the following, we focus on the modulations of enzymatic activities where the forward and backward reactions cannot be modulated independently ^3^. However, when the forward and backward reactions are independently modulated, for example by controlling the external concentrations of chemicals, we can extend our method to deal with such situations by treating the forward and backward reactions as independent reactions.

Note that the popular (bio)chemical reaction kinetics have the form shown in Eq, (8)-(10), such as the mass-action kinetics, (generalized) Michaelis-Menten kinetics, ordered- and random many-body reaction kinetics, and the ping-pong kinetics [44]. An important feature of the reaction kinetics of this form is that the direction of the reaction is set by the thermodynamic part as ***p***(***x***) is related to the thermodynamic force of the reaction. On the other hand, the remaining part ***f*** (***x***) is purely kinetic, and it modulates only the absolute value of the reaction rate, but not the direction. For reversible Michaelis-Menten kinetics, v_max_([S] − k[P])/(1 + [S]/K_*S*_ + [P]/K_*P*_), the thermodynamic part corresponds to its numerator, [S] − k[P], and the remaining part v_max_/(1 + [S]/K_*S*_ + [P]/K_*P*_) is the kinetic part^4^.

Now, we allow the enzyme activities ***u*** to temporally vary in 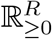 and compute the controllable set of a given target state ***x***\*, C(***x***\*). The model equation is given by

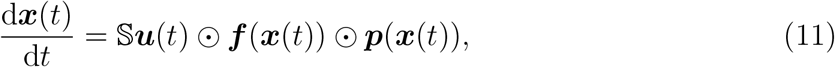

where ⊙ denotes the element-wise product.

First, we divide the phase space 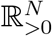 into subsets based on the directionalities of the reactions. Note that the kinetic part is strictly positive and 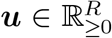 holds. Consequently, the directionalities of the reactions are fully set only by the thermodynamic part ***p***(***x***) ^5^. Let ***σ*** be a binary vector in {1, −1}^*R*^, and define the *direction subset* W_***σ***_ as

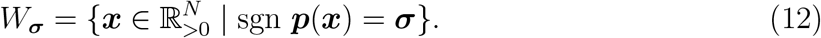

Next, we introduce the *balance manifold* of reaction *r* given by

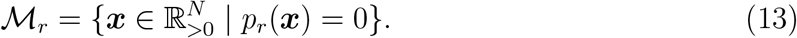

Now the phase space 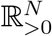 is compartmentalized by the balance manifolds 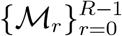 into the direction subsets W_***σ***_^6^.

Let *path* ***ξ***(*t*) be the solution of Eq. (11) with a given ***u***(*t*) reaching the *target state* ***x***\* at t = T from the *source state* ***x***_0_, i.e., ***ξ***(*T*) = ***x***\* and ***ξ***(0) = ***x***_0_, meaning that ***x***_0_ ∈ *C*(***x***\*). Additionally, we assume that if ***ξ***(*t*) intersects with any ℳ_*i*_, the intersection is transversal ^7^. As ***ξ***(*t*) is the solution of Eq. (11), ***ξ***(*t*) is given by

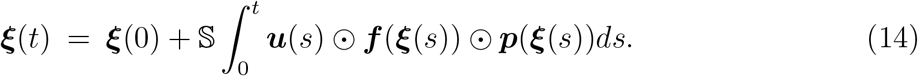

Next, we fragment the path. Note that the union of all the closures of the direction subsets, 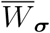, covers 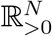, and ***ξ***(*t*) intersects transversally to the balance manifolds, if any. Thus, we can divide the interval [0, *T*] into *L* + 1 segments *I*_*i*_ = (*τ*_*i*_, *τ*_*i*+1_), (0 ≤ *i* ≤ *L*) so that 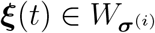 holds for *t* ∈ *I*_*i*_. In each interval *I*_*i*_, Eq, (14) is simplified as

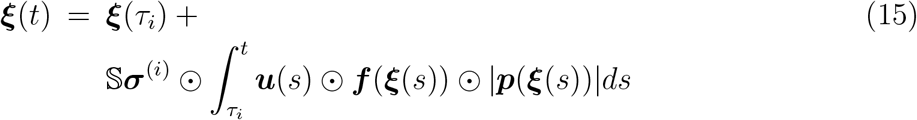

Recall that all the functions inside the integral are non-negative. Thus, ***ũ*** (*t*) := ***u***(*t*) ⊙ ***f*** (***ξ***(*t*)) ⊙ |***p***(***ξ***(*t*))| satisfies 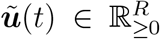. This means that we can consider the ramped function ***ũ*** (t) as a control parameter. By introducing 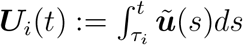, Eq, (15) is further simplified as

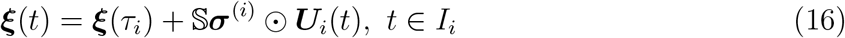

Therefore, for the solution of Eq.(11), ***ξ***(*t*) with ***ξ***(0) = ***x***_0_ and ***ξ***(*T*) = ***x***\*, the following holds; There exist a set of time-intervals 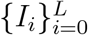, reaction directionalities 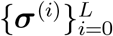, and non-negative-valued functions 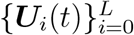 such that path in the interval *t* ∈ *I*_*i*_ is represented in the form of Eq, (16).

The converse of the above argument holds. Suppose that there is a continuous path 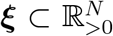 with ***x***_0_ and ***x***\* as its endpoints. Since ***ξ*** is a continuous path, it is continuously parameterizable by *t* ∈ [0, *T*] so that ***ξ***(0) = ***x***_0_ and ***ξ***(*T*) = ***x***\* hold. By following the above argument conversely, we can see that if there exist 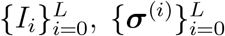, and 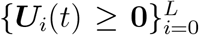 satisfying Eq, (16) and d***U***_*i*_(*t*)/dt ≥ **0**, there exists a control ***u***(*t*) given by

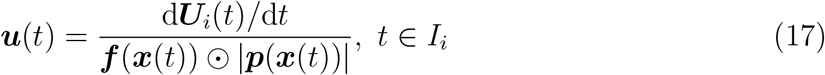

and 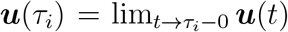, where the division is performed element-wise. With this ***u***(t), ***ξ***(t) satisfies Eq, (11). Thus, ***x***_0_ is an element of the controllable set C(***x***\*).

The summary of the above conditions leads to the definition of the *single stoichiometric path*.

### Definition 4

(Single Stoichiometric Path). *A continuously parameterized path* 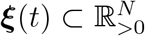 *is called a single stoichiometric path from* ***x***_0_ *to* ***x***\* *with signs* {***σ***^(*i*)^}^*L*^ *if there exists*

1. 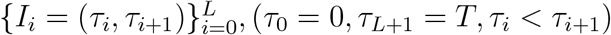, (*τ*_0_ = 0, *τ*_*L*+1_ = *T, τ*_*i*_ < *τ*_*i*+1_),
2. 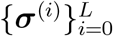, (sgn ***p***(***x***_0_) = ***σ***^(0)^, sgn ***p***(***x***\*) = ***σ***^(*L*)^, ***σ***^(*i*)^ ≠ ***σ***^(*i*+1)^),
3. 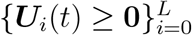

*satisfying the following conditions;*

1. ***ξ***(0) = ***x***_0_ *and* ***ξ***(*T*) = ***x***\*.
2. 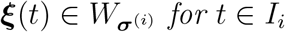 *for t* ∈ *I*_*i*_.
3. ***ξ***(*t*) = ***ξ***(*τ*_*i*_) + 𝕊***σ***^(*i*)^ ⊙ ***U***_*i*_(*t*) *for t* ∈ *I*_*i*_.
4. d***U***_*i*_(*t*)/d*t* ≥ **0** *for t* ∈ *I*_*i*_.

Let SP_*L*_(***x***\*) denotes the source points of all the single stoichiometric paths to ***x***\* with all possible choices of the sign sequence with a length less than or equal to L, and SP(***x***\*) := lim_*L*→∞_ SP_*L*_(***x***\*). We call SP(***x***\*) *the stoichiometric paths* while SP_*L*_(***x***\*) is termed *the finite stoichiometric paths* with length L. Note that the stoichiometric paths equals to the controllable set.

The stoichiometric paths is a useful equivalent of the controllable set, whereas the conditions 2 and 4 in the definition are laborious to check whether a given path and ***U***_*i*_(*t*) satisfies them. Thus, we introduce *the single stoichiometric ray*. It is easy to compute, and the collection of them gives exactly the stoichiometric paths if the thermodynamic part ***p***(***x***) of a model equation is linear.

Let us show that if ***p***(***x***) is linear, the single stoichiometric ray guarantees the existence of the single stoichiometric path. Since ***p***(***x***) is linear, all W_***σ***_’s are either convex polytopes or polyhedra. Suppose there is a set of points

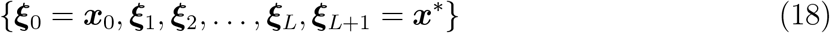

Satisfying

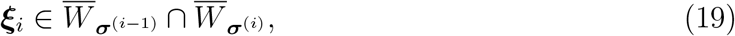

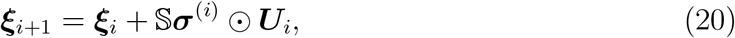

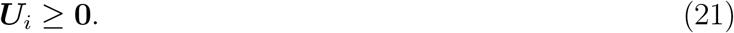

Then, we can construct a line graph ***ξ***(*t*) parameterized by *t* ∈ [0, *T*] which connects the points 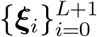 as

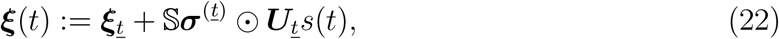

where

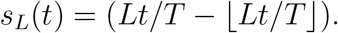

*t* is defined as ⌊t/(*LT*)⌋ with ⌊·⌋ as the floor function. Note that the function *s*_*L*_(*t*) is a repetition of the of linear function from 0 to 1 for *L* times (see Fig. 2 for graphical representations). Here, ***U***_*t*_s_*L*_(*t*) ≥ **0** and d(***U***_*t*_s_*L*_(*t*))/d*t* ≥ **0** follow. Additionally, the points between ***ξ***_*i*_ and ***ξ***_*i*+1_ are in the direction subset 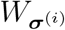. Thus, ***ξ***(*t*) and ***U***_*i*_ satisfy the conditions 2 to 4 in Def. 4 in *t* ∈ *I*_*i*_ = (*iT*/*L*, (*i*+1)*T*/*L*,). Also, since we choose 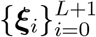 so that ***ξ***_0_ = ***x***_0_ and ***ξ***_*L*+1_ = ***x***\*, the line graph represented by Eq. (22) satisfies all the conditions in Def. 4.

**FIG. 2.**
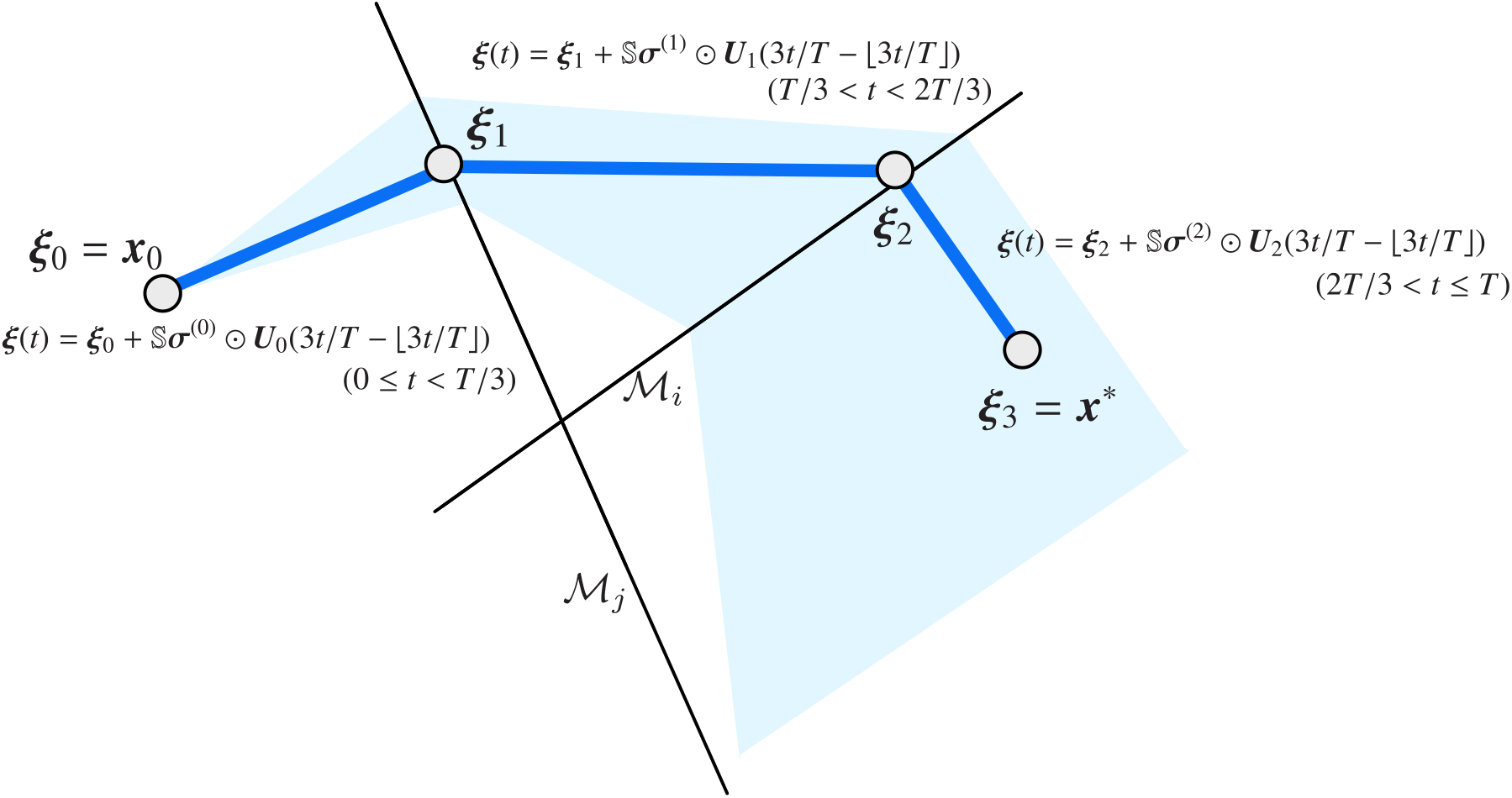
A visualisation of a single stoichiometric ray (blue line graph) defined in Eq. (22). Since the function *s*_*L*_(*t*) = (*Lt/T* − ⌊*Lt/T* ⌋) is a repetition of the linear function from 0 to 1 for *L* times, Eq. (22) represents a line graph connecting the vertices 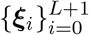. ℳ_*i*_ and ℳ_*j*_ are the balance manifolds of reaction *i* and *j* respectively. Cyan shaded region is the region reachable from ***x***_0_ by a single stoichiometric path.

We call this line graph *the single stoichiometric ray*. If there is a single stoichiometric ray from ***x***_0_ to ***x***\*, then there is a single stoichiometric path from ***x***_0_ to ***x***\*. Thus ***x***_0_ is an element of the stoichiometric paths SP(***x***\*). Note that this is only true if ***p***(***x***) is linear.

Let us formally introduce the single stoichiometric ray.

### Definition 5

(Single Stoichiometric Ray). *A line graph* ***ξ***(*t*) *parameterized by t* ∈ [0, *T*] *satisfying the following conditions is called a single stoichiometric ray from* ***x***_0_ *to* ***x***\* *with signs* 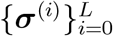 *where* sgn ***p***(***x***_0_) = ***σ***^(0)^, sgn ***p***(***x***\*) = ***σ***^(*L*)^, ***σ***^(*i*)^ ≠ ***σ***^(*i*+1)^.

1. *τ*_0_ = 0 < *τ*_1_ < · · · < *τ*_*L*_ < *τ*_*L*+1_ = *T*,
2. ***ξ***(0) = ***x***_0_ *and* ***ξ***(*T*) = ***x***\*,
3. 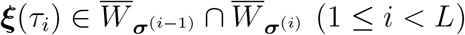,
4. ***ξ***(*τ*_*i*_ + *s*(*τ*_*i*+1_ − *τ*_*i*_)) = ***ξ***(*τ*_*i*_) + 𝕊***σ***^(*i*)^ ⊙ ***U***_*i*_s, (*s* ∈ [0, 1), ***U***_*i*_ ≥ **0**).

Let us define SR_*L*_(***x***\*) and SR(***x***\*), termed *the finite stoichiometric rays with length* L and the *stoichiometric rays* ^8^ in the same manner as the stoichiometric path. Note that only the evaluations of L discrete points are required to determine the existence of a single stoichiometric ray.

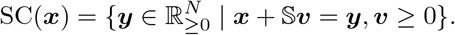

Indeed, if all the reactions of the model are irreversible, the stoichiometric rays of x* is equivalent to the stoichiometric cone with the replacement of 𝕊v by − 𝕊v.

Let us remark that SR_*L*_(***x***\*) = SP_*L*_(***x***\*) holds if the thermodynamic part ***p***(***x***) is linear, and thus, the stoichiometric rays is equivalent to the controllable set. If the thermodynamic part is nonlinear, SP_*L*_(***x***\*) ⊆ SR_*L*_(***x***\*) holds. If a single stoichiometric path connecting ***x***_0_ to ***x***\* exists, corresponding single stoichiometric ray exists which is the line graph connecting from ***x***_0_ to ***x***\* via the intersection points between the stoichiometric path and the balance manifolds (see Fig. 3A). However, the converse does not hold. Note that the stoichiometric path checks the consistency of the reaction direction at all points on the path, while the stoichiometric ray checks the consistency only at the discrete points on the manifolds (Fig. 3B). Thus, the stoichiometric rays may overestimate the controllable set, while the computational cost is much lower than the stoichiometric paths.

**FIG. 3.**
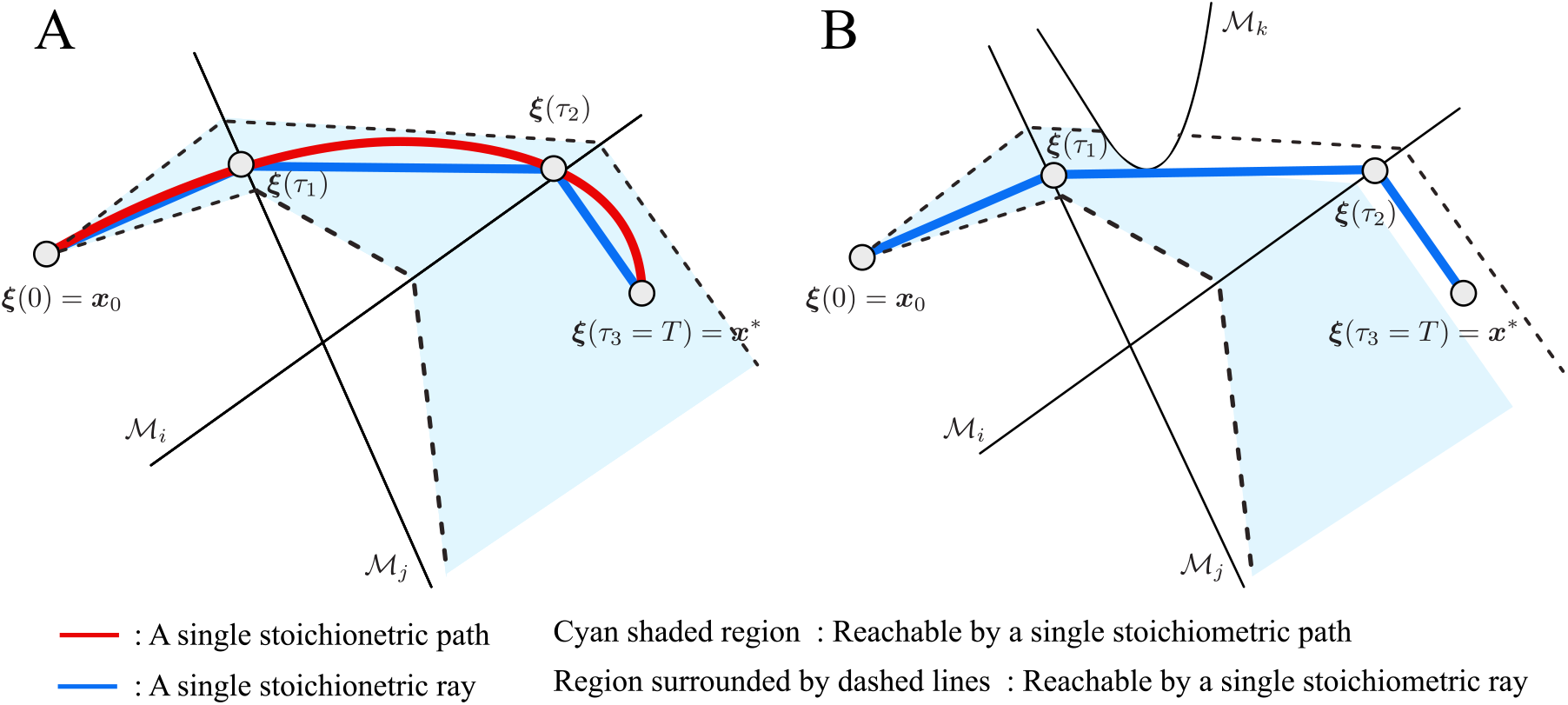
(A) A single stoichiometric path (red) and a single stoichiometric ray (blue) from ***x***_0_ to ***x***\* with 2 reaction direction flips. The cyan shaded region is the reachable region from ***x***_0_. The single stoichiometric ray is an approximation of the single stoichiometric path with a line graph. ℳ_*i*_ and ℳ_*j*_ are the balance manifolds of reaction *i* and *j*, respectively. (B) A case where there is no single stoichiometric path from ***x***_0_ to ***x***\*, but there is a single stoichiometric ray. The cyan shaded region is reduced by the blocking by the balance manifold ℳ_*k*_ (here we assume that the transition through the interior of ℳ_*k*_ is not possible), and thus the stoichiometric paths from ***x***_0_ to ***x***\* do not exist. However, since the stoichiometric rays evaluate the existence of the states at the boundary of the direction subsets, ignoring whether the paths are blocked inside the direction subset, a single stoichiometric ray is judged to exist. The region surrounded by the dashed lines is the reachable region by a single stoichiometric ray. Note that not only the single stoichiometric path shown in red in panel A is blocked, but also all paths from ***x***_0_ to ***x***\*.

Let us now show how the stoichiometric rays is computed using the reversible Brusselator model as an illustrative example. The model equation is given by

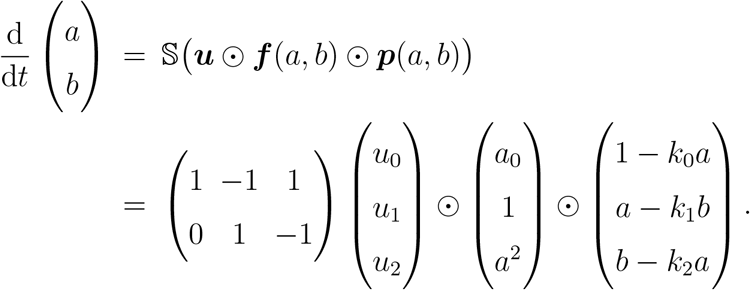

Note that the thermodynamic part vector ***p*** in the reversible Brusselator is a linear function of a and b whereas the reaction rate function vector ***f*** ⊙ ***p*** is a nonlinear function.

The finite stoichiometric rays for the two choices of the target state ***x***\* are shown in Fig. 4 (For the computational procedure, see SI text section S1 and SI Codes). In Fig. 4, SR_*L*_(***x***\*)\SR_*L*−1_(***x***\*) is filled with distinct colors. As shown in Fig 4A, it is possible to reach the point ***x***\* = (0.6, 0.6) from anywhere in the phase space, 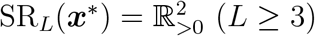, while we need to choose the starting point from the specific region for the other target point ***x***\* = (1.5, 1.5) as in Fig. 4B because 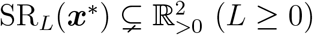 holds for the point.

**FIG. 4.**
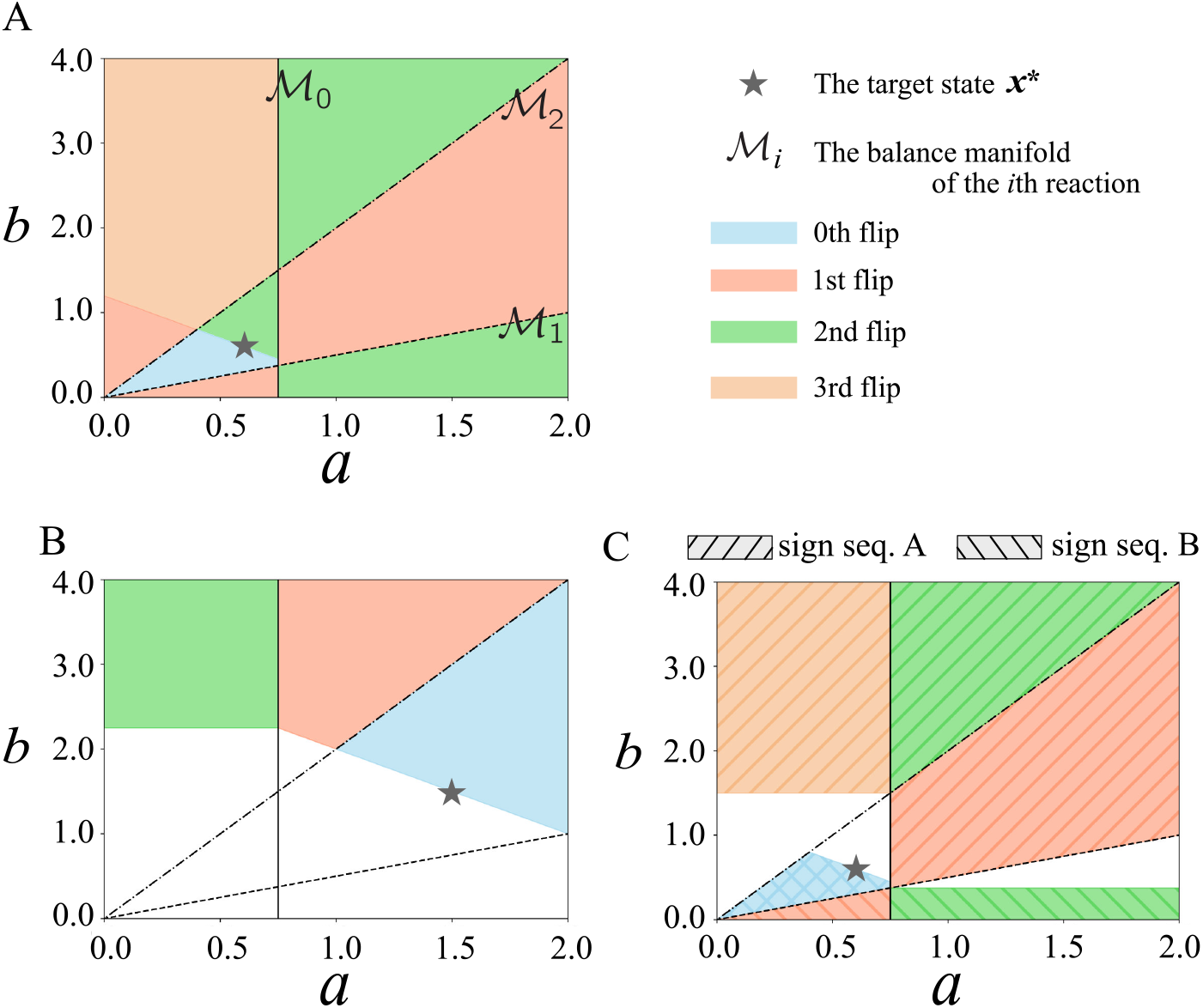
The stoichiometric rays of the reversible Brusselator model. The balance manifold ℳ_0_, *ℳ*_1_, and ℳ_2_ are represented by the solid, dash, and dot-dash lines, respectively. The phase space is restricted to [0, 2] × [0, 4] for computation. (A) The stoichiometric rays of ***x***\* = (0.6, 0.6). The regions covered for the first time at each flip of the reaction direction are filled with different colors. The whole region is covered within 3 flips. (B) The stoichiometric rays of ***x***\* = (1.5, 1.5). In this case, the bottom left region is never covered even if we allow arbitrarily number of flips. (C) The unions of all single stoichiometric rays of ***x***\* = (0.6, 0.6) with two example sign sequences are depicted. The sign sequence A and B follow are (1, −1, −1) → (−1, −1, −1) → (−1, −1, 1) → (1, −1, 1) and (1, −1, −1) → (1, 1, −1) → (−1, 1, −1), respectively. ***k*** = (4*/*3, 2, 2) is used.

## IV. DEATH OF A TOY METABOLIC MODEL

### A. Computing the dead set

Next, we apply the stoichiometric rays to a toy model of metabolism and compute its dead set. The model is an abstraction of the glycolytic pathway consisting of the metabolites X, Y, ATP, and ADP (a network schematic and reaction list are shown in Fig. 5A).

**FIG. 5.**
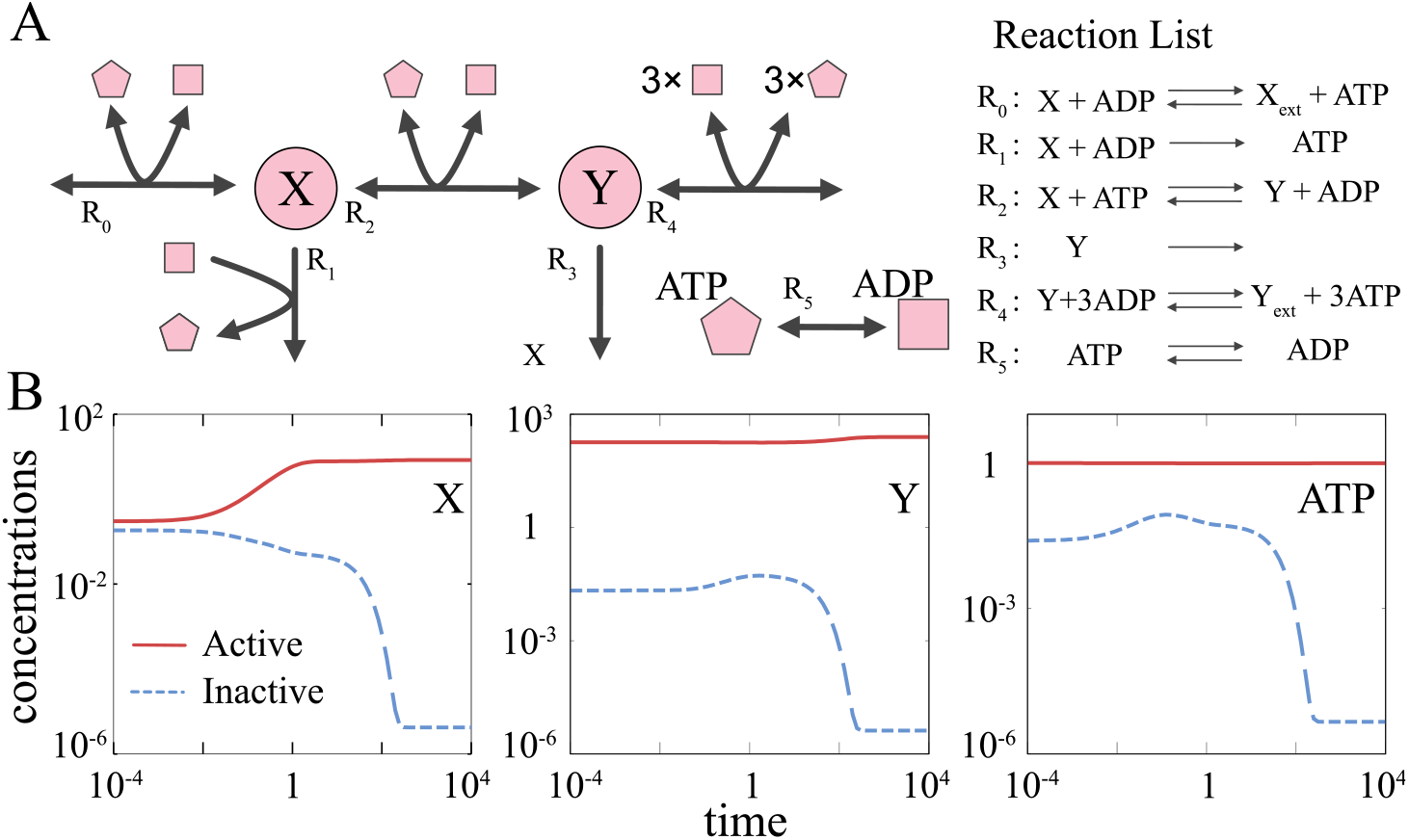
(A) A schematic of the reaction network. The list of the reactions is shown on the right side of the diagram. (B) An example of dynamics of metabolites X, Y, and ATP converging to the active (red), and inactive attractors (blue), respectively. Parameters are set to ***u*** = (1.0, 10.0, 0.1, 0.01, 1.0, 0.1) and *k*_0_ = 10.0, *k*_2_ = 0.1, *k*_4_ = 1, *k*_5_ = 10^−5^.

The rate equation is given by ^9^

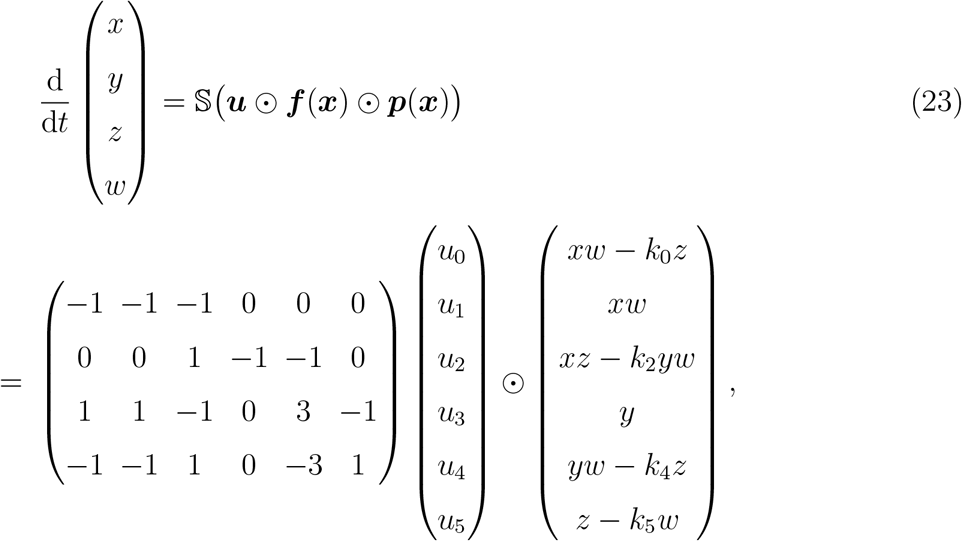

where x, y, z, and w represent the concentrations of metabolites X, Y, ATP, and ADP, respectively. In the second line of the above equation, ***f*** (***x***) = **1** is omitted. The thermodynamic part of the model is nonlinear. As the total concentration of ATP and ADP is a conserved quantity in the model, we replace w by *z*_*tot*_ − *z* where *z*_*tot*_ is the total concentration of the two (we set *z*_*tot*_ to unity in the following).

As shown in Fig. 5B, the model exhibits bistability with a given constant ***u***. On one attractor (the endpoint of the red line), X molecules taken up from the environment are converted into Y and secreted into the external environment resulting in the net production of a single ATP molecule per single X molecule. In contrast at the other attractor (the endpoint of the blue line), almost all X molecules taken up are secreted to the external environment via reaction R_1_. Y is also taken up from the external environment while consuming 3ATPs. Note that reaction R_0_ consumes a single ATP molecule to take up a single X molecule, and the reaction R_1_ produces a single ATP molecule. For the model to obtain a net gain of ATP, X must be converted into Y by consuming one more ATP molecule via reaction R_2_. However, once the model falls into the attractor where ATP is depleted (endpoint of the blue line in Fig. 5B), it cannot invest ATP via R_2_, and the model is “dead-locked” in the state. Hereafter, we refer to the former and latter attractors as the active attractor ***x***^*A*^ and inactive attractor ***x***^*I*^, respectively.

Now, we compute the dead states of the model according to Def. 3. In this framework, we first need to select the representative living states. Here, we suppose that the active attractor ***x***^*A*^ is the representative living state. According to Def. 3, we compute the complement set of the stoichiometric paths of the active attractor, 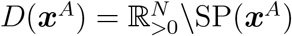. This complement set corresponds to the dead set with ***x***^*A*^ as the representative living state. Note that choosing ***x***^*A*^ as the representative living state is equivalent to choosing mutually reachable attractors from/to ***x***^*A*^ by continuously changing the enzyme activities and their basin of attraction (Recall Eq. (A.11)). By selecting a single-point attractor as the representative living state, we can effectively select a large region in the phase space.

Due to the challenges associated with computing the stoichiometric paths, we utilize the stoichiometric rays for the computation. Let us term 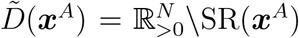 the *underestimated dead set*. As the stoichiometric rays is a superset of the stoichiometric paths, the underestimated dead set is a subset of the dead set. Thus the stoichiometric rays does not show a false negative of returnability; If 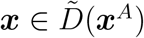 holds, then ***x*** is guaranteed to be dead with respect to the active attractor.

In Fig. 6A, the underestimated dead set with a finite *L*, 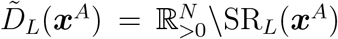 is depicted in the phase space (For the computational procedure, see SI text section S3 and SI Codes). The points inside the underestimated dead set cannot return to the active attractor regardless of how the enzyme activities are modulated with a given number of possible flips. The inactive attractor is contained in the underestimated dead set, and thus the inactive attractor is indeed the “death” attractor at least with the *L* = 4 flips ^10^. Also, the boundary of the overestimated controllable set 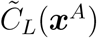 and the underestimated dead set 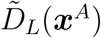 now gives an estimate of the SANZ surface, 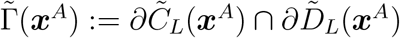^11^.

**FIG. 6.**
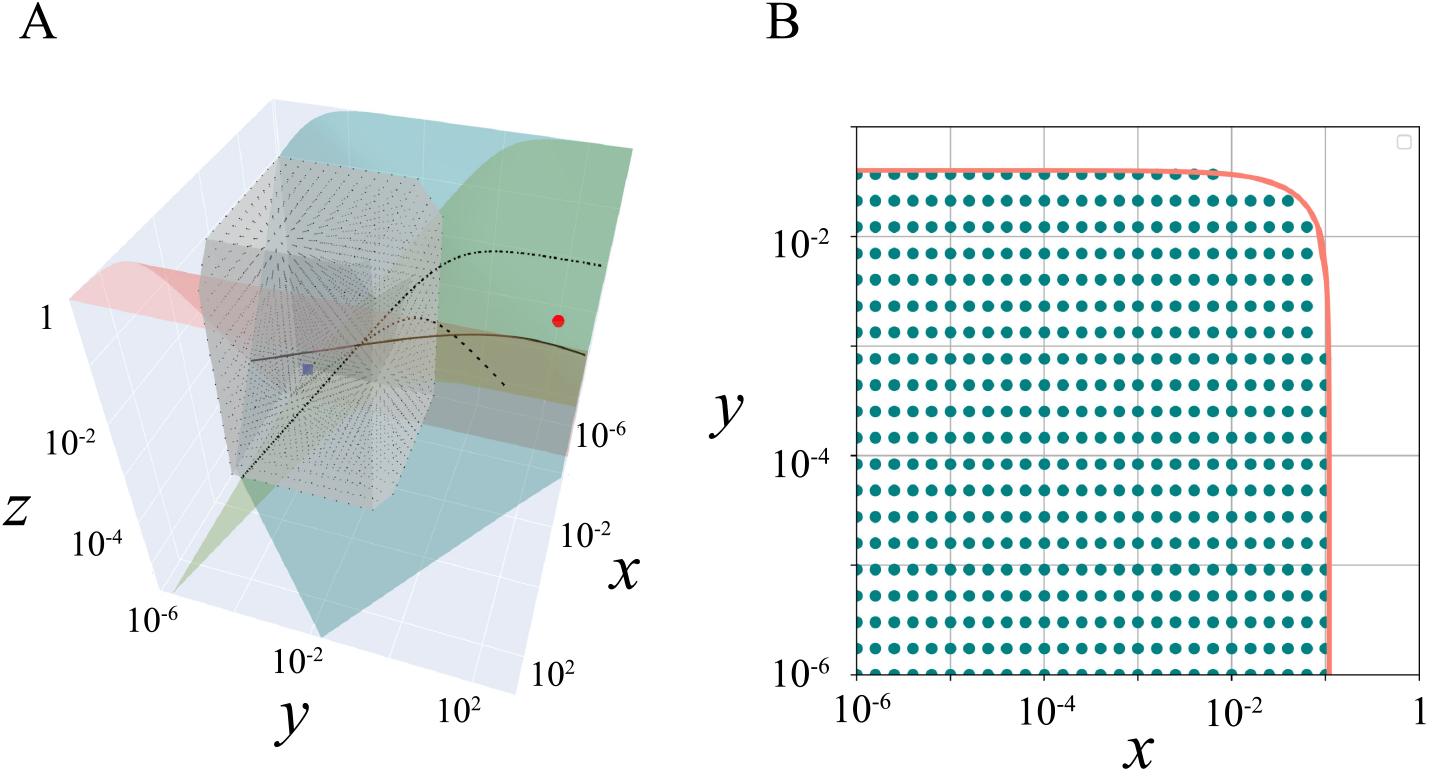
(A) The active attractor (red point), the inactive attractor (blue box), and the points in the underestimated dead set (black dots) are plotted. The gray rectangle is the convex hull of the dead states. The balance manifolds ℳ_0_, ℳ_2_, and ℳ_4_ are drawn as the salmon, green-blue, and olive-colored surfaces. The balance manifold ℳ_5_ is not illustrated because it is not necessary for the argument. The intersections of the balance manifold pairs (ℳ_0_, ℳ_2_), (ℳ_2_, ℳ_4_), and (ℳ_4_, ℳ_0_) are represented by the solid, dash, and dot-dash curves, respectively. (B) An analytic estimate of the boundary of the underestimated dead set (red curve) and the points in the underestimated dead set (green-blue points) are plotted. log_10_ *z* value is fixed to −1.7. We set *L* = 4. The same ***k*** values with Fig. 5 are used.

In Fig. 6B, we plot the points in the underestimated dead set with an analytically estimated SANZ surface of the set on the 2-dimensional slice of the phase space where z value is set to a single value. The SANZ surface is calculated from the thermodynamic and stoichiometric constraints that each single stoichiometric ray should satisfy. The curves match well with the boundary of the underestimated dead set (for the detailed calculation, see SI Text section S4).

### B. The global transitivity of the toy metabolic model

The application of the stoichiometric rays is not limited only to computations of the dead set with pre-defined representative living states. The stoichiometric rays enables us to investigate the mutual transitivity among states without a priori definition of living states. In this section, we present a transition diagram of the toy metabolic model ^12^. The transition diagram thus dictates the global transitivity of the model. This provides insights for choosing the representative living states, and in addition, criteria for the model selection suitable for investigating cell death.

For the computation of the transition diagram (procedure of the computation is illustrated in Fig. 7A), we sampled several points in the phase space (Fig. 7B) in addition to the active and inactive attractors, and computed controllability for all pairs of the sampled points. Note that we computed only the controllability to the active attractor in the previous part, but here we compute the controllability of all pairs of points mutually. With the computation, we construct the directed graph *G*_*L*_ where the nodes are the sampled points and the directed edges represent the controllability from the source point to the target point, i.e, we put the directed edge from ***x*** to ***y*** if ***x*** ∈ SR_*L*_(***y***) holds.

**FIG. 7.**
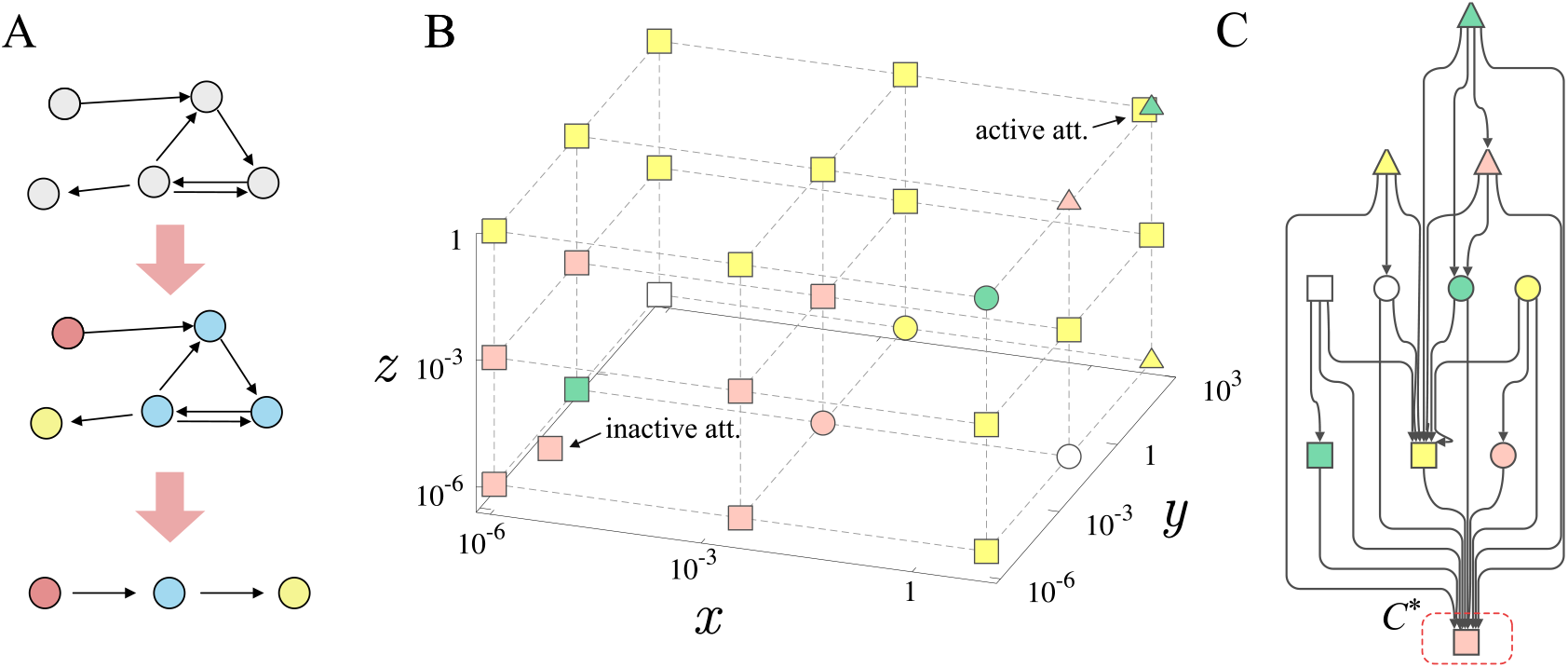
The transition diagram of the discrete states of the toy metabolic model. (A) Construction of the transition diagram. First, the controllability from/to all pairs of points are computed and the directed graph is drawn (top graph). In the graph, the directed edge represents that there are controls from the state at the tail of the arrow to the state at the head of the arrow. Then the nodes are grouped into strongly connected components (middle graph). The component graph is obtained by merging the nodes belonging to the same strongly connected components into a single node (bottom graph). (B) The points for which the mutual transitivity is computed. The points are grouped and labeled by the strongly connected components (SCCs). The dashed lines are eye guides. The active attractor (yellow square) and the inactive attractor (pink square) are shown together. Note that the green triangle is on the grid, but not the active attractor. (C) The transition diagram of the SCCs. Only the two SCCs (pink- and yellow squares) contain more than one state. The terminal SCC (pink square, indicated by the red-dashed rectangle), *C*\*, has only in-edges. The same ***k*** values with Fig. 5 are used, and *L* = 4.

Next, we computed the condensation of *G*_*L*_ which, denoted 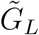. Condensation is a directed graph in which strongly connected components (SCCs) of *G*_*L*_ are contracted to a single node. The nodes of *G*_*L*_ within the same SCC are mutually reachable ^13^. Hereafter, we refer to the condensation graph 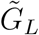 as a transition diagram.

The transition diagram is shown in Fig. 7C ^14^. In addition, because the condensation graph is a directed acyclic graph, the diagram highlights the hierarchy of the state transitions or the “potential” of the states. The active and inactive attractors are in the SCCs represented by the yellow- and pink squares in Fig. 7C, respectively. Let us refer to the pink SCC as the terminal SCC, C*, since it has only in-edges, and thus, the states in 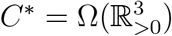 (cf. Def. 9 in Appendix) cannot escape from the set regardless of the control ***u***(t).

The computation of the transition diagram provides a guide for selecting the representative living states, and in addition, a criterion for the model selection suitable for investigating cell death. It is commonly believed that controlling to the living states is difficult and possible only from limited states. Also, all living and non-living systems can be non-living (dead state), i.e., control to the dead state should always be possible regardless of the source state. If we accept these beliefs, the representative living state should be taken from the SCCs except for the terminal SCC. When we additionally require that the representative living state be the attractor (i.e., the state is stable without temporal modulation of enzymatic activities), the representative living state should be taken from the SCC containing the active attractor (the yellow square in Fig. 7C).

In the toy metabolic model, the active attractor and the inactive attractors are in the non-terminal SCC and the terminal SCC, respectively. However, this is not always the case for any biochemical model. In order to study cell death employing mathematical models, we may require the models to satisfy conditions such as the one above. The transition diagram is useful method for checking if a given model satisfies requirements as the diagram captures the global transitivity in the model.

## V. DISCUSSION

Questions such as “what is life?” and “what is death?” relate to how we can characterize living and dead states. Providing a straightforward answer for such a characterization is challenging even for mathematical models of cells. However, quantitative evaluation of the “life-death boundary” (SANZ hypersurface), or in other words, characterization of the difference between life and death can be possible. In the present manuscript, we proposed a theoretical framework to study the dead set and SANZ hypersurface with the development of a tool for computing it.

In the present framework, the dead state is defined based on the controllability to the representative living state. Thanks to the useful mathematical features of controllability relationship, ***x*** ⇀ ***y***, the death judgment is robust on the choice of the representative living state; The choice of representative living state is arbitrary, as long as they are chosen from the same equivalence class. Further, Thm. 1 (see Appendix) states that even if there are uncountably many living states, the representative living states can be countably many, or even, a finite number.

Our strategy for defining the dead state is to use the law of exclusion, of the form “what is not living is dead”. This allows us to avoid direct characterization of dead states. In this sense, our characterization of death is similar to that of the regrow experiment, rather than the staining assay in which the dead state is characterized by stainability. For such indirect characterizations, it is necessary to determine whether a cell will eventually regrow. This is an extremely hard challenge in experimental systems. However, by focusing only on mathematical models, we can determine, in principle, whether the cell will regrow. The stoichiometric rays provides a reasonable method for computing this.

Note that the present framework says nothing about which states should be chosen as representative living states. We assume that information about which states should be considered “living” is given by other sources, such as experiments and observations. The present framework is intended to provide a quantitative criterion for the irreversible transition from the living states to the dead states with respect to a given representative living states. Yet, the living/dead judgment is not that sensitive to the choice of representative living states. We have shown that choosing several states as the representative living states is typically equivalent to choosing a large variety of states as the representative living states (see Thm. 1 in the Appendix). Also, if the dead set with respect to the first choice of representative living states X has states that are “obviously living” by other criteria, one can update the representative living states to X^′^ by adding those states to X and compute the dead set D(X^′^). The present framework does not define life and death as if it were a transcendent being, but rather computes the dead states and the SANZ hypersurface based on one’s working hypothesis of “what is living”.

The heart of the stoichiometric rays is that the modulation of enzymatic activity cannot directly change the directionalities of the reactions. As a consequence of this feature, the balance manifolds and direction subsets are invariant to enzymatic activities. This allows us to efficiently compute the stoichiometric rays. The usefulness of the stoichiometric rays was demonstrated by using two simple nonlinear models of chemical reactions. Especially, in the toy metabolic model, we showed that the stoichiometric rays is a powerful tool for computing the dead set in which no control can return the state to the active attractor with a given number of flips.

Thermodynamic (ir)reversibility is believed to be key to grasping the biological (ir)reversibility such as the irreversibility of some diseases and death. However, there is no system-level link between thermodynamic and biological reversibilities except in a few cases where simple causality leads to biological irreversibility such as in transmissible spongiform encephalopathy [47]. The stoichiometric rays enables us to bridge the (ir)reversibility of each reaction to the system-level reversibility; Recall that the SANZ surface of the dead set is calculated from the thermodynamic constraint, i.e., directionality of the reactions, and the stoichiometric constraint. Further studies should pave the way to unveil the relationship between thermodynamic irreversibility and biological irreversibility. This may reveal the energy (or “negentropy” [48]) required to maintain cellular states away from the SANZ hypersurface [49].

We do not intend to extend the scope of the framework indiscriminately, while it is noteworthy that the framework does not assume any specific mechanism of cell death. There are multiple cell death mechanisms such as apoptosis, necrosis, ferroptosis, pyroptosis, autophagy, erebosis, and so on [50–55]. The molecular details and cellular behaviors during death processes differ depending on the mechanism. However, the irreversible transition to a state where multiple cellular functions are lost is common to all mechanisms. Our framework provides a quantitative criterion for the irreversible transition, and thus its applicability is not limited to specific mechanisms.

Finally, we would like to mention an extension of the present framework to the probabilistic formulation and its implications. In the present deterministic version, the transition from the living state to the dead state is irreversible. In stochastic processes, however, the transition from the state considered dead in the deterministic model to the living states is possible with some probability. Thus, in probabilistic formulation, life and death are not binary categories, but are continuously linked states. Given the small size of cells, the probabilistic formulation is more suitable for describing cellular state transitions. Death is commonly believed to be irreversible. However, if the theory of death reveals the probabilistic nature of life-death transitions, it may have multiple implications for reforming the concept of “death”.

## Appendix The choice of the representative living states

In this section we make some statements about the choice of representative living states. The main claim is that controllability leads to the equivalence relation and the partial order. The equivalence relation shows that there are pairs of states such that choosing one of them as the representative living state results in the same dead set as choosing both of them. The partial order shows that there is a reduction of the representative living state without changing the dead set.

Let us start introducing the the controllability relationship.

### Definition 6

(⇀ and ⇌). *We denote* ***x*** ⇀ ***y*** *if* ***x***, 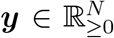 *satisfy* ***x*** ∈ C(***y***). *If* ***x*** ⇀ ***y*** *and* ***y*** ⇀ ***x*** *holds, we denote* ***x*** ⇌ ***y***.

### Proposition 1.

⇀ *is preorder and* ⇌ *is equivalence relation on* 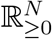.

The proof is straightforward from the definition of the controllable set.

From the proposition, we obtain the following corollary.

### Corollary 1.

*For any* ***x***, 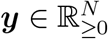 *satisfying* ***x*** ⇀ ***y***,

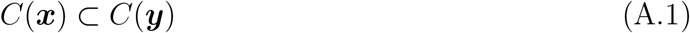

*holds. If* ***x*** ⇌ ***y*** *holds, we have*

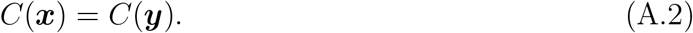

*Proof*. For any ***z*** ∈ *C*(***x***), ***z*** ⇀ ***x*** holds. From the assumption, we also have ***x*** ⇀ ***y***. Thus, the transitivity of ⇀ results in ***z*** ⇀ ***y***. Therefore, *C*(***x***) ⊂ *C*(***y***) holds. By repeating the same argument for ***y*** ⇀ ***x***, we have the second statement.

Thereby, if the states ***x***, 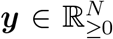 are mutually controllable to each other ***x*** ⇌ ***y***, then the dead set is the same whether ***x*** or ***y*** is chosen as the representative living state. This implies that we can use the quotient set 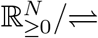 for studying the controllable set of the states instead of the original space 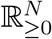.

### Definition 7

(Quotient set and Equivalence class). *The quatient set of* 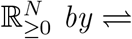 *is repre-sented by* 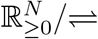, *and the equivalence class of* ***x*** *is denoted by* 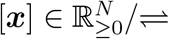.

### Definition 8

(↬). *For any* [***x***], 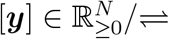, *if* ***x*** ∈ [***x***] *and* ***y*** ∈ [***y***] *exist such that* ***x*** ⇀ ***y*** *holds (see Def. 6), we denote* [***x***] ↬ [***y***].

### Proposition 2.

↬ *is partial order on* 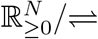.

*Proof*. Note that if we have [***x***] ↬ [***y***], ***x*** ⇀ ***y*** holds for any ***x*** ∈ [***x***] and ***y*** ∈ [***y***].

Reflexibility: Since ***x*** ⇀ ***x*** holds, [***x***] ↬ [***x***] holds.

Transitivity: If [***x***] ↬ [***y***] and [***y***] ↬ [***z***] hold, ***x*** ⇀ ***y*** and ***y*** ⇀ ***z*** hold. Thus, ***x*** ⇀ ***z***

holds from the transitivity of ⇀, and [***x***] ↬ [***z***] holds.

Antisymmetry: If [***x***] ↬ [***y***] and [***y***] ↬ [***x***] hold, ***x*** ⇀ ***y*** and ***y*** ⇀ ***x*** hold. Thus, ***x*** ⇌ ***y***

holds, and we have [***x***] = [***y***].

### Corollary 2.

*For any* [***x***], 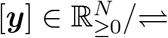, *if* [***x***] ↬ [***y***] *holds, we have*

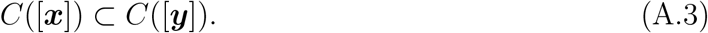

*Proof*. The controllable set of the equivalence class [***x***] is a union of all points in [***x***], i.e., 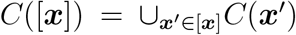. From Cor. 1, the controllable set of every ***x*** ∈ [***x***] is identical. Therefore, we can take a representative state ***x*** ∈ [***x***] and C([***x***]) = C(***x***) holds. The same argument for [***y***] leads to C([***y***]) = C(***y***), (***y*** ∈ [***y***]). By appliying Cor. 1 to C(***x***) and C(***y***), we obtain C([***x***]) = C(***x***) ⊂ C(***y***) = C([***y***]).

This corollary states that the “larger” equivalence class in terms of ↬ ^15^ has the larger controllable set. This motivates us to introduce the *terminal class* of the representative living states X.

### Definition 9

(Terminal Class). *For a given set* 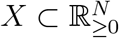, *we take the set of the equivalence classes* 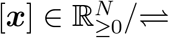 *with nonempty intersection with* X *and denote it by* 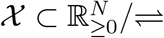 *i*.*e*.,

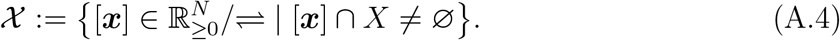

*We call the maximal element of* 𝒳 *with respect to* ↬ *the terminal class of X. We denote the set of all terminal classes of X by* Ω(*X*) *(for* 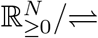 *and* ↬, *see Def. 7 and Def. 8, respectively*.).

Note that an equivalence class [***x***\*] ∈ 𝒳 being the terminal class means that there is no other equivalence class [***x***] ∈ 𝒳, ([***x***] ≠ [***x***\*]) satisfying [***x***\*] ↬ [***x***] exists.

Ω(*X*) can be an empty, finite, or infinite set. However, if we can assume that every chain in 𝒳 (a totally ordered subset of 𝒳) has an upper bound with respect to ↬, Zorn’s lemma guarantees Ω(X) being nonempty. To satisfy this condition, it is sufficient that the model (Eq. (1)) has a choice of the constant control parameter ***u*** with which the model never shows a divergent behavior.

The controllable set of each terminal class [***x***\*] contains that of any other equivalence class satisfying [***x***] ↬ [***x***\*]. This allows us to reduce the representative living states X into a smaller set X* without changing the dead set.

For 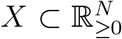, suppose that Ω(*X*) is nonempty. Let us take a point from each terminal class in Ω(*X*) and denote the set of the chosen points by 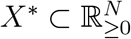, that is,

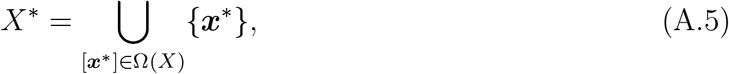

where the choise of the representative element from [***x***\*] is arbitrary. We have the following theorem for the dead set with respect to *X* and *X*\* (For a graphical representation of the theorem, see Fig. 8).

**FIG. 8.**
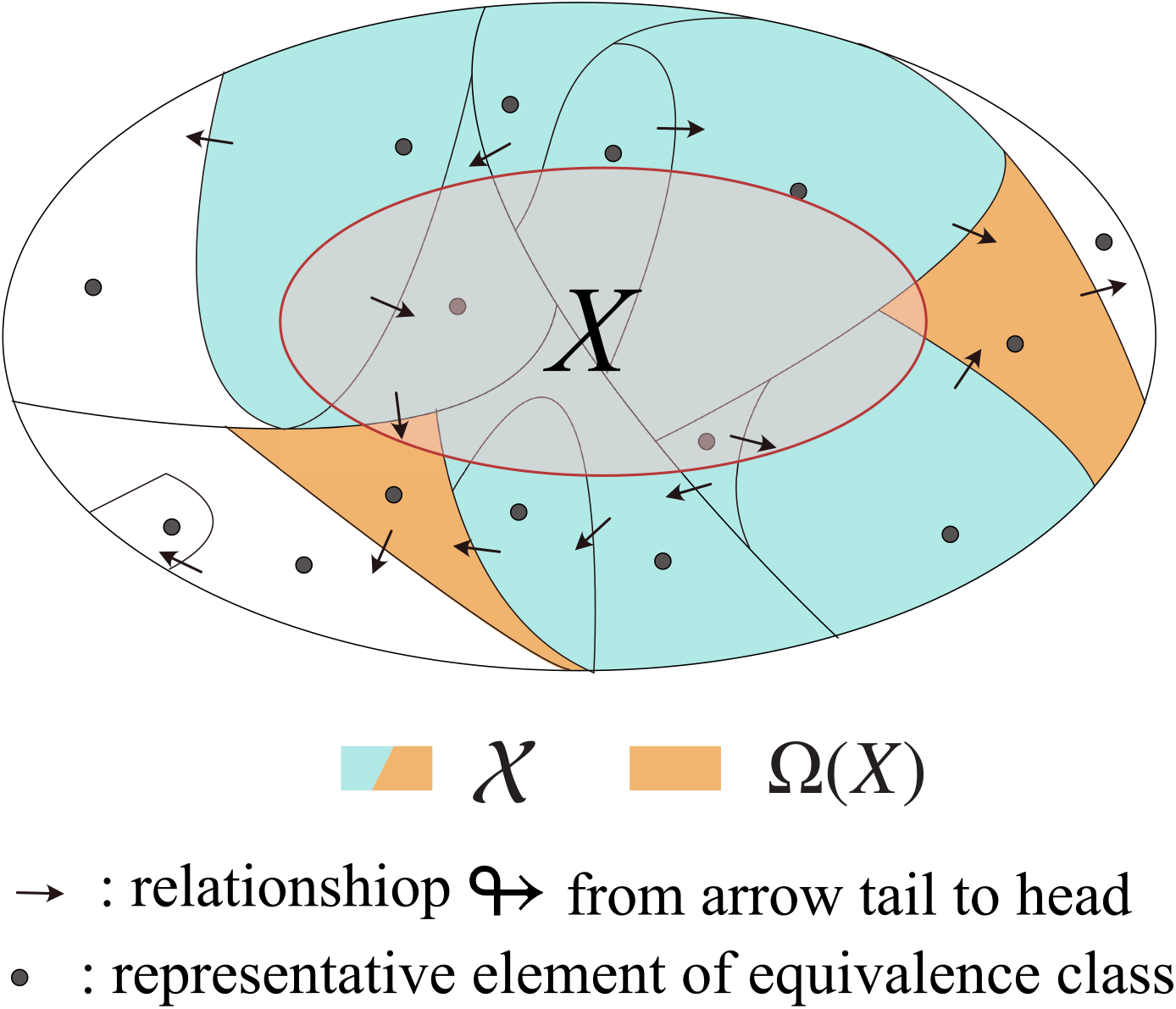
The equivalance classes and the representative living states *X*. The large and small ellipses represent 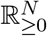 and the representative living states *X*, respectively. Each compartment is the equivalence class 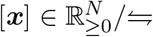. Where the arrow represents the partial order ↬ from the tail of the arrow to the head. Each black point is the representative element of each equivalence class. The set X is the set of equivalence classes filled with cyan or orange. The terminal classes Ω(*X*) ⊂ 𝒳 is the set of equivalence classes filled with orange. According to Theorem. 1, the controllability to *X* is fully captured by computing the controllability to the representative elements of [***x***] ∈ Ω(*X*).

### Theorem 1.

*The dead set with respect to X is identical to the dead set with respect to* X*, *i*.*e*.,

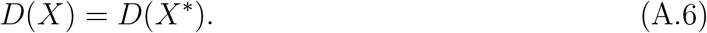

*Proof*. Recall that *D*(*X*) = *D*(*X*\*) is equivalent to *C*(*X*) = *C*(*X*\*). The controllable set of *X* is defined as

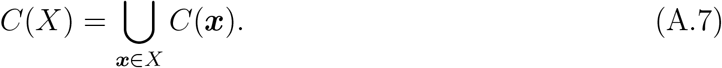

As the controllable set of the equivalence class *C*([***x***]) and the controllable set of a point ***x*** ∈ [***x***] are identical (Cor. 1), we have

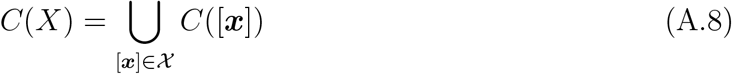

where 𝒳 is what defined in Def. 9.

For each [***x***] ∈ 𝒳, if there exists [***y***] ∈ 𝒳, ([***y***] ≠ [***x***]) such that [***y***] ↬ [***x***] holds, we have *C*([***y***]) ⊂ *C*([***x***]) from Cor. 2. Therefore, we can drop *C*([***y***]) from the union in Eq. (A.8) without violating the equality. By repeating the dropping process, only the terminal classes remain and we obtain the following equality:

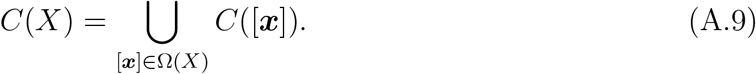

Additionally, since *C*([***x***]) = *C*(***x***) holds for any representative point ***x*** ∈ [***x***], we have

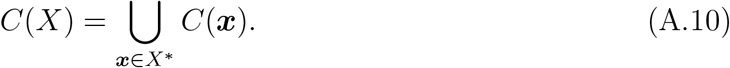

The theorem states a useful property for choosing the representative living states to compute a dead set. Suppose that we have the representative living state 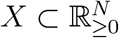. Without the theorem, we must check the controllability of all points in *X* to judge the dead state. However, with this theorem, we can take discrete points from each terminal class in Ω(*X*) and check the controllability of the chosen points to judge the dead state. Note that the set *X*\* can also be finite, which enables further efficient computations of the dead set.

An example of practical applications of this theorem is as follows. Suppose that the model has a point attractor ***a***_***u***_ with a fixed control parameter value ***u***, and ***a***_***u***_ is a continuous function of ***u*** in *U* ⊂ ℝ^*P*^ . In such cases, taking the point attractor with a single parameter choice ***u***_0_ ∈ *U* is equivalent to taking the basin of attraction *B*(***a***_***u***_) for all ***a***_***u***_, as the representative living states, i.e.,

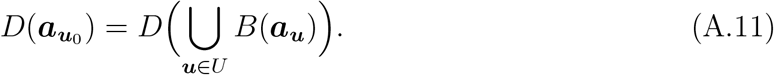

This is because 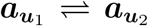 holds for any ***u***_1_, ***u***_2_ ∈ *U* and also we have ***x*** ⇀ ***a***_***u***_ for any ***x*** ∈ *B*(***a***_***u***_).

## ACKNOWLEDGMENTS

We thank Chikara Furusawa and Ikumi Kobayashi for the discussions. This work is supported by JSPS KAKENHI (Grant Numbers JP22K15069, JP22H05403 to Y.H.; 24K00542, 19H05799 to T.J.K.), JST (Grant Numbers JPMJCR2011, JPMJCR1927 to T.J.K.), and GteX Program Japan Grant Number JPMJGX23B4. S.A.H. is financially supported by the JSPS Research Fellowship Grant No. JP21J21415.

## SI CODES

All codes for the main result are deposited on GitHub (https://github.com/yhimeoka/StoichiometricRay).

## Footnotes

1 An intuitive terminology is to call the state in *D*(*X*) with no temporal change (d***x****/dt* = 0) “dead” (i.e. at the fixed point) and “dying” otherwise. However, in the present framework, whether a given steta is dead or dying depends on the specific value of the control parameter. Moreover, one can make all states in *D*(*X*) to be dying (d***x****/*d*t* ≠ 0) states by keep changing the control parameter, for example by setting ***u***(*t*) = ***u***_0_| sin(*t*)|, where ***u***_0_ is a constant vector. Thus, in the present framework, the distinction between dying and dead is neither essential nor possible.

2 In typical cases, the method works for 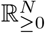, including the two examples presented in the latter half of the manuscript. However, it is known that special care is necessary for mathematical claims at the boundary (*x*_*i*_ = 0 for one or more chemicals) [43]. In the present manuscript, we avoid dealing with the boundary.

3 The enzymes change the activation energy of reactions to facilitate the reaction rate, but cannot directly change the chemical equilibrium, i.e. the direction of the reactions [42].

4 The separation of ***f*** (***x***) and ***p***(***x***) is not necessary. However, the separation is useful to distinguish between the thermodynamic contribution and the kinetic contribution to the reaction rate, i.e. ***f*** depends on the enzymatic reaction mechanism (it is whether Michaelis-Menten, ping-pong, mass-action, etc.), whereas ***p*** does not. The separation also simplifies the calculation of the balance manifold, which will be introduced later.

5 For the directionality of the *r*th reaction at *u*_*r*_ = 0, we define it by sgn 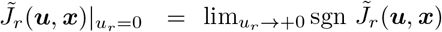

6 On the balance manifolds, 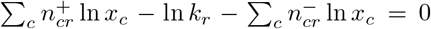 ln *x*_*c*_ = 0 holds. This is a linear equation of ln ***x***, and thus, each direction subset is connected. Depending on how the balance manifolds intersect, the number of the direction subset changes. At most, there are 2^*R*^ direction subsets for a model with *R* reactions.

7 In the followings, we assume the transversality of the intersections between ***ξ***(*t*) and ℳ_*i*_.

8 The term “stoichiometric rays” is derived from the stoichiometric cone. The stoichiometric rays is a generalized concept of the stoichiometric cone [43] defined by

9 For a derivation of the linear thermodynamic part for the reaction R_4_, see SI text section S2.

10 We computed the underestimated dead set for *L* = 6 with a lower resolution of the point sampling due to the computational cost. We confirmed that the increase of *L* from 4 to 6 does not make the underestimated non-returnable set larger. (see Fig. S3).

11 The non-emptiness of the dead set is due to the fact that negative-valued controls are not allowed. It is shown from the rank relation of the Lie algebra of the model that if negative values are allowed for the control parameters to take, controls from any state to any state are feasible [45, 46]. In particular, the model shows a bifurcation at ***u*** ≈ (1.0, 10.0, 0.1, −0.08, 1.0, 0.1), where the inactive attractor becomes unstable, and the model shows the unistability of the active attractor. Thus, modulating *u*_3_ to ≈ −0.08 is a way to revive it.

12 The computation of the transition diagram corresponds to the computation of the equivalent classes and partial order (↬, 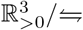) defined in the appendix section.

13 Each node in the condensation graph 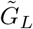 corresponds to an equivalent class (see Appendix).

14 The transition diagram describes the partial order relation ↬ on 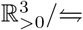 (see Appendix).

15 Here, we say that [***x***] is “larger” than [***y***] if [***y***] ↬ [***x***] holds.

## Notes

### Competing Interest Statement

The authors have declared no competing interest.

### Summary of Updates

According to the referees' comments of the journal, we reorganized the manuscript and added several new figures to increase the readability for non-math background readers.

