## Supplemental Information Text for "A theoretical basis for cell deaths"

### Supplementary Information Text of
### *A theoretical basis for cell deaths*

Yusuke Himeoka, Shuhei A. Horiguchi, and Testuya J. Kobayashi

### S1 Computational procedure of the controllable set for the model with linear thermodynamic part

**Algorithm S1** Algorithm for computing the stoichiometric rays with a given sign sequence

**Input:** $\{\boldsymbol{\sigma}^{(1)}, \boldsymbol{\sigma}^{(2)}, \cdots, \boldsymbol{\sigma}^{(L)}\}, \boldsymbol{x}^{(\mathrm{tgt})}$

*Initialisation* :

$X \leftarrow \{\boldsymbol{x}^{(\mathrm{tgt})}\}$

$\mathcal{R} \leftarrow \{\boldsymbol{x}^{(\mathrm{tgt})}\}$

*LOOP Process*

**for** $i = 1$ to $L$ **do**

  $X_{\mathrm{ex}} \leftarrow$ `ExPt_Enumerate`$(X, \boldsymbol{\sigma}^{(i)})$

  $\mathcal{R} \leftarrow \mathcal{R} \cup$ `ConvexHull`$(X_{\mathrm{ex}})$

  **if** $i < L$ **then**

    $r \leftarrow$ `Diff`$(\boldsymbol{\sigma}^{(i)}, \boldsymbol{\sigma}^{(i+1)})$

    $X \leftarrow X_{\mathrm{ex}} \cap \mathcal{M}_r$

  **end if**

**end for**

**return** $\mathcal{R}$

In `ExPt_Enumerate`, the extreme points of the polytope are enumerated. The function `ConvexHull` returns the convex hull of the given set of points. The function `Diff` returns the reaction index having the opposite sign of the two sign vectors (In the algorithm, we handle the case where the reaction flips one by one).

In `ExPt_Enumerate`, we compute the extreme points of the polytope given by

$$P(X, \boldsymbol{\sigma}) = \{\boldsymbol{y} \in \Omega \mid \boldsymbol{y} = \sum_{\boldsymbol{x} \in X} \lambda_{\boldsymbol{x}} \boldsymbol{x} - \mathbb{S}\boldsymbol{\sigma} \odot \boldsymbol{v}, \boldsymbol{v} \geq \boldsymbol{0}, \lambda_{\boldsymbol{x}} \geq 0, \sum_{\boldsymbol{x} \in X} \lambda_{\boldsymbol{x}} = 1\} \cap W_{\boldsymbol{\sigma}},$$

where $\Omega$ is the region for the numerical computation. The above polytope is the reachable points from the points in $X$ (for the first loop, $\boldsymbol{x}^{(\mathrm{tgt})}$) by the backward time evolution, and thus, from any point in the polytope, points in $X$ are reachable by the forward time evolution.

The extreme points enumeration is executed using the python package `pypoman` [1] which computes the extreme points of the polytope represented by the linear inequality $A\boldsymbol{y} \leq \boldsymbol{b}$ (For details, see supplemental code). The function `ConvexHull` is implemented using `scipy.spatial.ConvexHull`. `Diff` function is a self-made function that returns the index of the reaction that has the opposite sign of the two sign vectors.

Using the algorithm.S1, we computed the finite stoichiometric rays with length $L$ by the following algorithm.

**Algorithm S2** Algorithm for computing the finite stoichiometric rays with the length $L$

**Input:** $L, \boldsymbol{x}^{(\mathrm{tgt})}$

*Initialisation* :

$SR_L \leftarrow \emptyset$

$\Sigma \leftarrow$ all possible sign sequences of length $L$

*LOOP Process*

**for** $i = 1$ to $|\Sigma|$ **do**

  $\mathcal{R} \leftarrow$ `Algorithm.S1`$(\Sigma_i, \boldsymbol{x}^{(\mathrm{tgt})})$

  $SR_L \leftarrow SR_L \cup \mathcal{R}$

**end for**

**return** $SR_L$

### S2 Derivation of the reaction rate function of multi-body reaction with the reaction degree 1

Here we present how a multi-body reaction

$$2X \rightarrow 2Y$$

can lead to the reaction rate function proportional to $[X]$, not $[X]^2$.

Suppose a reaction scheme shown in Fig.S1. The reaction is composed of three steps; the formation of the enzyme-substrate complex $[EX]$, irreversible maturation of the enzyme-substrate complex for the next step, the formation of the enzyme-substrate complex $[E^*XX]$, and the formation of the products. The reaction rate constants are denoted by $k^a_+, k^a_-, k^b_+, k^c_+, k^c_-, v$. The temporal evolution of the concentrations is given by the following equations

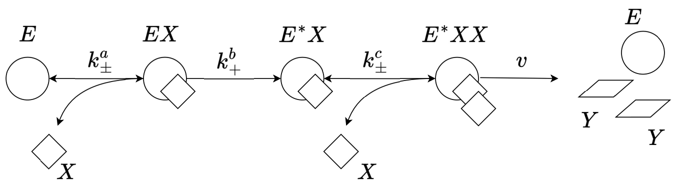


Figure S1: The schematics of the enzymatic reaction $2X \rightarrow 2Y$.

$$\frac{\mathrm{d}[EX]}{\mathrm{d}t} = k_+^a[E][X] - k_-^a[EX] - k_+^b[EX] \quad \text{(S1)}$$

$$\frac{\mathrm{d}[E^*X]}{\mathrm{d}t} = k_+^b[EX] - k_+^c[E^*X][X] + k_-^c[E^*XX] \quad \text{(S2)}$$

$$\frac{\mathrm{d}[E^*XX]}{\mathrm{d}t} = k_+^c[E^*X][X] - k_-^c[E^*XX] - v[E^*XX] \quad \text{(S3)}$$

The steady-state solution is given by

$$[EX] = \frac{k_+^a}{k_-^a + k_+^b}[E][X] \quad \text{(S4)}$$

$$[E^*X] = \frac{k_+^b}{k_+^c - k_+^c k_-^c/(v + k_-^c)} \frac{k_+^a}{k_-^a + k_+^b}[E] \quad \text{(S5)}$$

$$[E^*XX] = \frac{k_+^a k_+^b}{v(k_-^a + k_+^b)}[E][X] \quad \text{(S6)}$$

Thus, the production rate $v[E^*XX]$ becomes linear of $[X]$. By introducing one more maturation step after the formation of $[E^*XX]$ and the formation of $[E^*XXX]$, we can obtain the same, linear dependency of the production rate on $[X]$ for $3X \to 3Y$ as well.

### S3 The transitivity computation of a given pair of the source and target point

We formulate the computational procedure of the transitivity from a given source point to a given target point using the following optimization problem

$$\begin{aligned}
\text{minimize} \quad & \zeta \\
\text{subject to} \quad & \sum_i n_{ir}^+ y_i^{(j)} - \sum_i n_{ir}^- y_i^{(j)} - \ln k_r = 0, \ (1 \le j < L) \\
& \boldsymbol{x}^{(j)} = \exp\left(\boldsymbol{y}^{(j)}\right), \ (1 \le j < L) \\
& \boldsymbol{x}^{(j)} + \mathbb{S}\boldsymbol{\sigma}^{(j)} \odot \boldsymbol{v}^{(j)} + \boldsymbol{\xi}^{(j)} = \boldsymbol{x}^{(j+1)}, \ (0 \le j < L) \\
& \boldsymbol{v}^{(j)} \ge \boldsymbol{0}, \ (0 \le j < L) \\
& \boldsymbol{y}^{(0)} = \ln \boldsymbol{x}^{(\mathrm{src})} \\
& \boldsymbol{y}^{(L)} = \ln \boldsymbol{x}^{(\mathrm{tgt})} \\
& \zeta \ge 0 \\
& |\xi_i^{(j)}| \le \zeta, \ (0 \le i < L)
\end{aligned}$$

Here, the variables are $\boldsymbol{y}^{(j)}, \boldsymbol{x}^{(j)}, \boldsymbol{v}^{(j)}, \boldsymbol{\xi}^{(j)}$, and $\zeta$. The index $j$ runs from 0 to $L$.

The first constraint correspoins to the points $\boldsymbol{x}^{(1)}, \boldsymbol{x}^{(2)}, \cdots, \boldsymbol{x}^{(L-1)}$ are on the balance manifolds. The logarithm conversion of the concentration $\boldsymbol{x}$, denoted by $\boldsymbol{y}$, is useful in terms of the numerical stability and also for the non-integer reaction degree $n_{ir}^{\pm}$. Instead, the second constraint is necessary for the consistency of $\boldsymbol{y} = \ln \boldsymbol{x}$.

The third and fourth constraints require the existence of the reaction flux vector with a given directionality $\boldsymbol{\sigma}^{(j)}$, with that the state $\boldsymbol{x}^{(j)}$ can transit to $\boldsymbol{x}^{(j+1)}$. In the third constraint, the slack variable $\boldsymbol{\xi}^{(j)}$ is introduced so that the optimization problem is always feasible. As all the slack variables should be zero if there is a single stoichiometric ray from the source to the target if the optimal value of $\zeta = \max_{ij} |\xi_i^{(j)}|$ is zero (practically, we adopted $10^{-9}$ as the threshold for the optimality), we judge that the source point is reachable to the target point with a given sign sequence $\boldsymbol{\sigma}$.

### S4 Estimate of the SANZ surface of the non-returnable set

Here we present the details of an analytic estimate of the SANZ (separating alive and non-life zone) surface of the underestimated non-returnable set $\tilde{\mathcal{D}}_L(\boldsymbol{x}^L)$ on the 2-dimensional slice at $\log_{10} z = -1.7$. The estimate is based on an observation of the single stoichiometric rays to the live attractor from the points outside of the underestimated non-returnable set. We generated a point close to, but outside of $\tilde{\mathcal{D}}_L(\boldsymbol{x}^L)$ and gradually brought the point to the SANZ surface along the $Y$ axis, with computing a single stoichiometric ray to the live attractor. As shown in Fig. S2A, the single stoichiometric ray first transits into the direction subset surrounded by the three balance manifolds, and climbs up to the live attractor. The closer the source state gets to the SANZ surface ($y$ gets smaller), the more its single stoichiometric ray enters the direction subset near the tip of the subset (see also Fig. S2B). The same behavior is observed when a point is generated away from the SANZ surface in the X direction and brings the point to the SANZ surface along the X axis.

This direction subset contains the live attractor, and the reaction signs are given by $\boldsymbol{\sigma}^* = (-1, 1, 1, 1, 1, 1)$. Importantly, all the standard basis with both signs $\pm \boldsymbol{e}_i,\ (i = x, y, z)$ can be generated by the positive linear combination of $\{\sigma_i^* \boldsymbol{S}_{*i}\}_{i=0}^{5}$, where $\boldsymbol{S}_{*i}$ is the $i$th column vector of the stoichiometric matrix. Thus, once the state enters this subset, the state can reach any point in the subset, and accordingly, the returnability to the live attractor is determined by whether the transition to this subset is possible. Let us term this subset the *target subset* in the following.

From the observation in Fig. S2, we hypothesize that a point becomes non-returnable when no single stoichiometric ray from the point can reach the tip of the target subset, $\boldsymbol{x}_c = (x_c, y_c, z_c)$. The source points shown in Fig. S2A belong to the direction subset with $\boldsymbol{\sigma}^{(Y)} = (-1, 1, -1, 1, 1, 1)$. Let us derive the conditions to reach the tip from a given source point.

Note that in this direction subset, the positive-valued standard basis for $x$ can be generated by the positive linear combination of the reaction stoichiometric vectors $\sigma_0^{(Y)}\boldsymbol{S}_{*0}$ and $\sigma_5^{(Y)}\boldsymbol{S}_{*5}$, and $x < x_c$ holds for $x \in W_{\boldsymbol{\sigma}^{(Y)}}$. Thus, the x value is tunable to $x_c$ independently from the other variables. On the other hand, only the negative-valued standard basis exists in the direction subset for $y$ and $z$. Thus, the necessary and sufficient condition to reach the tip is as follows; there exists a point reachable from the source point by a combination of the reactions that are not generated by the positive linear combinations of the negative-signed standard basis, and each element of the point is larger than the corresponding element of the tip, i.e.,

$$y - v_2 - v_4 \geq y_c,\ z + v_0 + v_2 + 3v_4 \geq z_c,\ \boldsymbol{v} \geq 0$$

This condition is equivalent to satisfying either one or both of the following inequalities;

$$y - v_2 \geq y_c,\ z + v_0 + v_2 \geq z_c, \boldsymbol{v} \geq 0$$

and

$$y + v_4 \geq y_c,\ z + v_0 - 3v_4 \geq z_c, \boldsymbol{v} \geq 0$$

The former inequalities are the condition for reaching the target subset using the reaction $\mathrm{R}_0$ and $\mathrm{R}_2$ and the latter are for the paths with $\mathrm{R}_0$ and $\mathrm{R}_4$. By tidying up the conditions, we obtain

$$-n\delta y \geq \delta z \tag{S7}$$

as a necessary condition to fulfill the inequalities where $\delta y$ and $\delta z$ denotes $y_c - y$ and $z_c - z$, respectively. $n$ depends on the combination of the reactions. $n = 2$ for the combination of $\mathrm{R}_0$ and $\mathrm{R}_2$ while is 3 for $\mathrm{R}_0$ and $\mathrm{R}_4$. The same argument is applicable also in the subset with the sign $\boldsymbol{\sigma}^{(X)} = (-1, 1, 1, 1, -1, 1)$ and we obtain

$$-n(\delta x + \delta y) \geq \delta z \tag{S8}$$

The Sanzu surface depicted in Fig.4B in the main text is the curve satisfying Eq. (S7) and (S8).

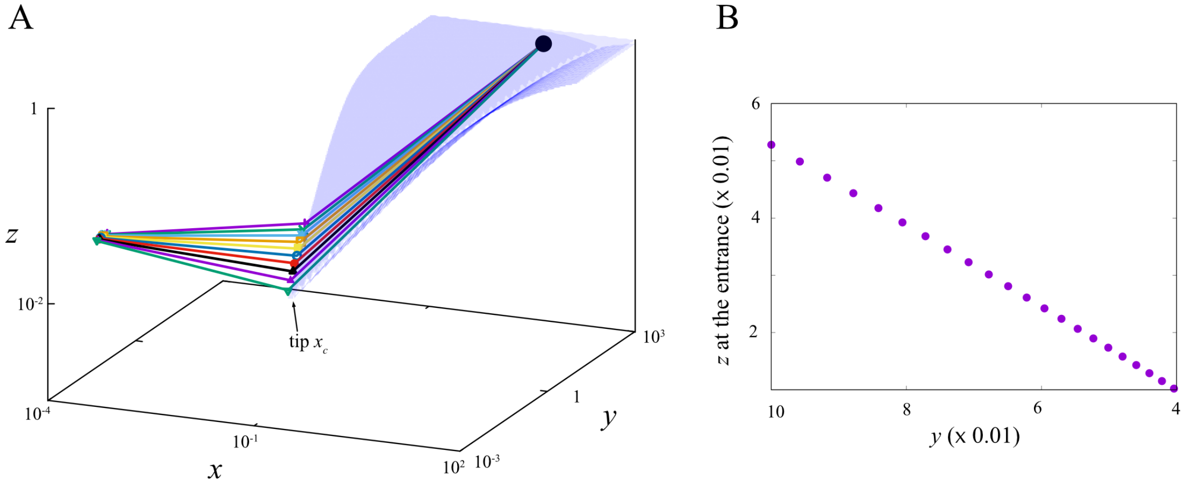


Figure S2: A. Single stoichiometric rays from the points close to the underestimated non-returnable set and the direction subset to which the active attractor belongs are depicted. As the source points approach the underestimated non-returnable set (small $y$ direction), the entrance point of the single stoichiometric ray becomes closer to the tip of the direction subset. B. The $z$ value at the entrance point is plotted as a function of the $y$ value of the source points. The $x$ and $z$ values of the source points are fixed to $10^{-4}$ and $10^{-1.7}$, respectively. We computed the single stoichiometric rays with the maximum possible $z$ value at the entrance point of the direction subset to clearly present the stoichiometric constraints.

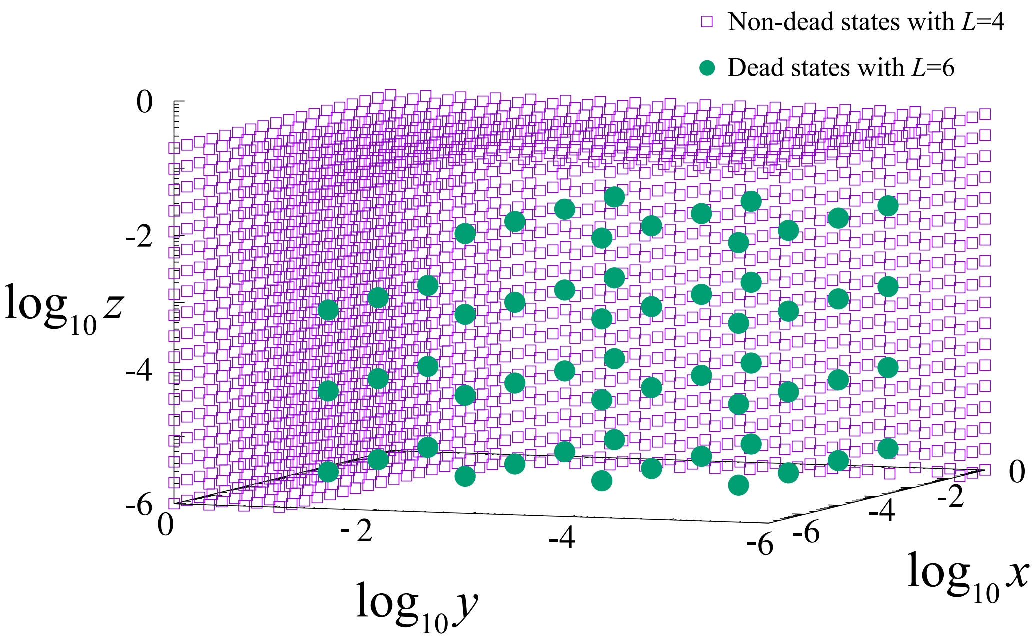


Figure S3: The dead states with $L = 6$ (filled green points) are plotted togeter with the non-dead states (states controllable to the active attractor $\boldsymbol{x}^A$) with $L = 4$ (open purple squares states). Five points were taken in each axis direction and the controllability were computed with the allowed maximum flips 6. No point overlaps with the non-dead states with $L = 4$, meaning that the increase of $L$ does not make the underestimated non-returnable set larger.
